# Long-term T cell perturbations and waning antibody levels in individuals needing hospitalization for COVID-19

**DOI:** 10.1101/2022.03.17.484640

**Authors:** Melissa Govender, Francis R. Hopkins, Robin Göransson, Cecilia Svanberg, Esaki M. Shankar, Maria Hjorth, Åsa Nilsdotter Augustinsson, Johanna Sjöwall, Sofia Nyström, Marie Larsson

## Abstract

COVID-19 is being extensively studied, and much remains unknown regarding the long-term consequences of the disease on immune cells. The different arms of the immune system are interlinked, with humoral responses and the production of high-affinity antibodies being largely dependent on T cell immunity. Here, we longitudinally explored the effect COVID-19 has on T cell populations and the virus-specific T cells, as well as neutralizing antibody responses, for 6-7 months following hospitalization. The CD8^+^ TEMRA and exhausted CD57^+^CD8^+^ T cells were markedly affected with elevated levels that lasted long into convalescence. Further, markers associated with T-cell activation were upregulated at the inclusion, and in the case of CD69^+^CD4^+^ T cells this lasted all through the study duration. The levels of T cells expressing negative immune checkpoint molecules were increased in COVID-19 patients and sustained for a prolonged duration following recovery. Within 2-3 weeks after symptom onset, all COVID-19 patients developed anti-nucleocapsid IgG and spike-neutralizing IgG as well as SARS-CoV-2-specific T cell responses. In addition, we found alterations in follicular T helper (TFH) cell populations, such as enhanced TFH-TH2 following recovery from COVID-19. Our study revealed significant and long-term alterations in T cell populations and key events associated with COVID-19 pathogenesis.

## Introduction

More than two years since initial reports of the disease, the COVID-19 pandemic has claimed approximately six million lives world-wide leaving survivors with long-term health consequences [1]. The etiologic agent, severe acute respiratory syndrome coronavirus 2 (SARS-CoV-2) gains entry into host cells after binding to angiotensin-converting enzyme 2 (ACE2) and transmembrane protease, serine 2 (TMPRSS2) via the receptor binding domain (RBD) of the viral spike (S) protein [2, 3]. While a vast majority of infected individuals remain asymptomatic or pauci-symptomatic, severe and life-threatening disease occurs in 14% and 5-6% of cases, respectively [4]. Approximately 20% of all infected individuals, especially the elderly and those with co-morbidities require hospitalization. In fatal cases of COVID-19, death often occurs due to organizing pneumonia (OP), acute respiratory distress syndrome (ARDS) or sepsis-like cytokine storm-associated multi-organ failure [5–7]. The hyperinflammation in COVID-19 is caused by the activated inflammatory cells recruited in response to infection and not by the virus per se [8–10]. Furthermore, the variations in initial host responses appear to determine the varied disease spectrum observed in COVID-19 [11, 12].

Clearance of viral infection involves both cellular and humoral immune responses [13, 14]. Virus-specific CD4^+^ and CD8^+^ T cells are present in almost all individuals who have had a prior episode of COVID-19 [15, 16], with lower numbers of these cells linked to disease severity [17–19]. Furthermore, the rapid appearance of functional SARS-CoV-2-specific T cells is linked to mild disease presentation [14]. The humoral immune response also plays a critical role in viral clearance in the host. The SARS-CoV-2-specific antibody responses vary in levels, as well as quality between asymptomatic individuals and those presenting with severe disease. While some studies have found higher levels and longer duration in more severe cases, others have reported no observable difference [20–26].

Follicular helper T cells (TFH) are primarily defined as CXCR5, programmed cell death protein 1 (PD-1), and Bcl-6 expressing CD4^+^ T cells. The frequency of circulating TFH cells is often low during homeostasis and is believed to mirror their numbers in the germinal centers of peripheral lymphoid organs, where they provide critical support to antigen-specific B cells to facilitate the development of long-term and high-affinity antibodies [27–31]. Effects on peripheral TFH cells have been documented in a few studies covering acute [32], mild [23], and severe/fatal SARS-CoV-2 infections [16]. However, not many have followed the infected population longitudinally [15, 33]. Peripheral TFH cells have been shown to be linked to neutralizing antibodies (nAb) in diseases such as HIV-1 [34], and now SARS-CoV-2 [22, 25, 35]. Furthermore, individuals who have recovered from severe compared to non-severe COVID-19 reportedly have higher nAb titers and CXCR3^+^ TFH cell frequencies, and faster recovery of lymphocyte levels, a month after leaving the hospital [36].

Viral infections lead to the activation of T cells with upregulated expression of markers that define early, i.e., CD69 and late, i.e., CD38/HLA-DR, activation [37, 38]. Activated CD4^+^ and CD8^+^ T cells exist during active/ongoing COVID-19 and in certain individuals, even after recovery, activated CD8^+^ T cells persist for over 100 days [39]. A case study of mild COVID-19 reported activated CD38^+^ HLA-DR^+^ CD8^+^ T cells at day 7, which surged to a peak at day 9 but remained high even at day 20. Activated CD4^+^ T cells were also present, but at lower levels [16]. Many studies support the activation of T cells and B cell responses during the onset of COVID-19 disease, but the infection also leads to transient immunosuppression, especially during severe disease [40]. The expression of negative immune checkpoint molecules on CD8^+^ and CD4^+^ T cells examined in 14 individuals with severe COVID-19 revealed an increased abundance of PD-1 and TIM-3 levels in most severe COVID-19 cases [17]. In addition, exhausted T cells expressing CD57 and PD-1, with impaired proliferation, were found in individuals with COVID-19 [41].

It is clear that COVID-19 induces immune dysfunction and alterations within T cell populations, and in light of this, we have longitudinally investigated the T cell profile of COVID-19 patients over a 6-7 month period, to monitor the changes in dynamics and functional quality of T cell responses. To further understand the immune responses elicited during SARS-CoV-2 infection, we also explored its long-term consequences on B cell and antibody responses in the cohort. We found an increase in TFH cells and alterations in TH1/TH2/TH17 TFH subsets that persisted following recovery from COVID-19, and a consistent rise in the CD8^+^ terminal effector memory RA^+^ (TEMRA) subset throughout the 6-7 months of the study. Further, markers associated with T-cell activation were upregulated and different forms of T cell impairment were also evident that lasted for a prolonged time following recovery from COVID-19.

## Materials and Methods

### Study Design

The present study was approved by the Swedish Ethical Review Authority (Ethics No. 2020-02580) and recruited hospitalized patients with COVID-19 (N=46; age range 32 - 91 years) and healthy staff (N=31; age range 23 - 62 years) at the Infectious Diseases Clinic at the Vrinnevi Hospital, Norrköping, Sweden. The participants were ≥18-years-old male and female subjects who provided written informed consent before enrollment. The study was carried out in compliance with good clinical practices, including the International Conference on Harmonization Guidelines and the Declaration of Helsinki. COVID-19 patients were enrolled into the study during hospitalization and first blood samples were drawn, which was followed by three additional samples collected at 2 weeks, 6 weeks, and 6-8 months, respectively after inclusion (**Figure 1**). The clinical and demographic characteristics of the hospitalized COVID-19 cohort are summarized in **Table 1**.

**Figure 1.**
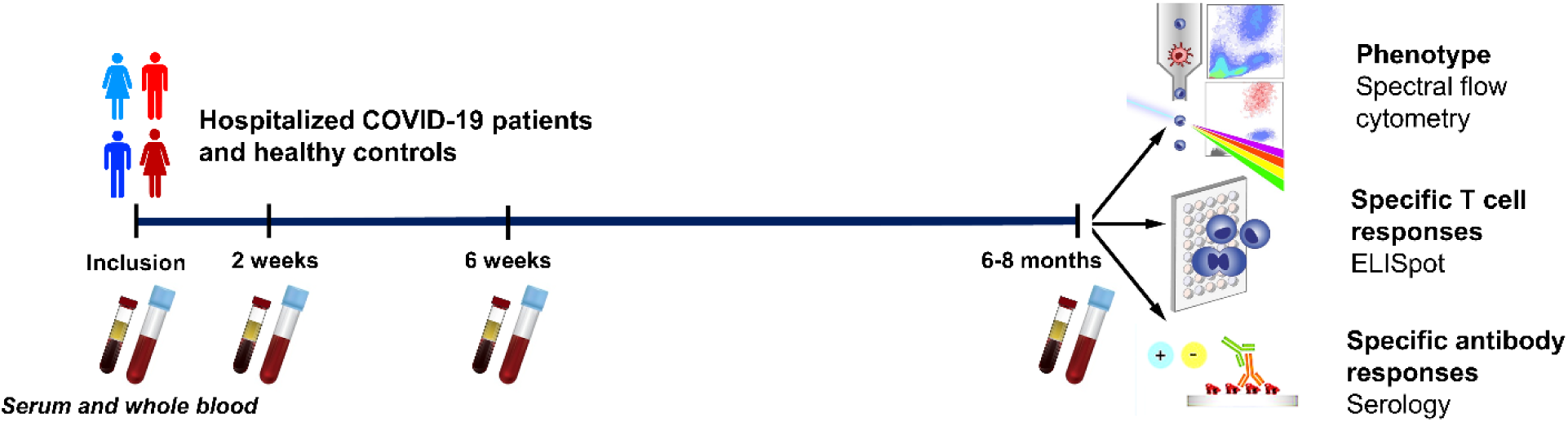
Graphical outline of the study. Peripheral blood was collected from patients with COVID-19 that needed hospitalization (N=46) and healthy controls (N=31). Clinical parameters were measured at enrollment, and the levels and effects on peripheral immune cells and neutralizing antibodies were assessed at enrollment, 2 weeks, 6 weeks, and 6-8 months. The phenotype of the T cells and B cells were analyzed by flow cytometry, SARS-CoV-2-specific T cells by ELISPOT, and SARS-CoV-2-specific antibodies by serological testing.

**Table 1.** Clinical and-demographical characteristics of hospitalized COVID-19 patients.

Diabetes mellitus, Cardiovascular, and Chronic pulmonary conditions defined as taking medication for these conditions
| Variable | Medical data | Reference range |
| --- | --- | --- |
| Number of patients | 46 |  |
| Age, median (range) | 56.6 (32-91) |  |
| BMI, median (range) | 30.2 (22.4-45.2) |  |
| Biological sex % | 34.8% F/65.2%M (16F/30M) |  |
| Days in hospital, median (range) | 8.8 (2-34) |  |
| ICU/ pandemic Ward %, (N) | 8.7/91.3 (4/42) |  |
| Days with symptoms before inclusion (median) | 9.5 (2-34) |  |
| Spike antibody IgG positive at inclusion %, (N) | 65.2 (30) |  |
| Nucleocapsid IgG positive at inclusion %, (N) | 63.0 (29) |  |
| Antiviral treatment %, (N) | 34.8 (16) |  |
| Corticosteroid treatment %, (N) | 67.4 (31) |  |
| No/oxygen/HFNOT:CPAP/mechanical ventilation % (N) | 6.5/41.3/50.0/2.2 (3/19/23/1) |  |
| Cardiovascular condition %, (N) | 56.5 (26) |  |
| Chronic pulmonary condition %, (N) | 23.9 (11) |  |
| Diabetes mellitus %, (N) | 26 (12) |  |
| Two of the underlying conditions %, (N) | 28.3 (13) |  |
| Three of the underlying conditions %, (N) | 4.4 (2) |  |
| Disease score: Moderate/severe %, (N) | 93.5 (43)/6.5 (3) |  |
| Smoker/snus %, (N) | 15.2 (7) |  |
| Previous history of smoking /snus %, (N) | 54.3 (25) |  |
| Leukocytes ($1 \times 10^9$/L) median (range) | 6.6 (2.8-20.4) | 3.5-8.8 |
| Platelets ($1 \times 10^9$/L) median (range) | 258 (121-543) | 150–400 |
| Lymphocytes ($1 \times 10^9$/L) median (range) | 1 (0.2-2.8) | 1.1-4.8 |
| Monocytes ($1 \times 10^9$/L) median (range) | 0.38 (0.1-1.24) | 0.1-1 |
| Lactate dehydrogenase (μKat/L) median (range) | 6.6 (3.2-16) | Over 70 <3.5 Under 70 < 4.3 |
| C-reactive Protein (mg/L) median (range) | 73 (6-318) | 0-10 |

### Patient Characteristics

On enrollment, the COVID-19 patients were provided with an approved questionnaire to secure details regarding COVID-19-associated symptoms prior to hospitalization, and smoking habits. Relevant medical data for the study were collected from digital medical records and included date of COVID-19 diagnosis, length of hospital stay, highest level of care received, maximum need of oxygen supplementation (<5L/min or HFNOT/CPAP/mechanical ventilation), details of COVID-19-related medication (anti-coagulants, remdesivir and/or dexamethasone), current medication, underlying conditions (if any) such as cardiovascular disease (CVD), chronic pulmonary disease, chronic renal failure, diabetes mellitus, and immunosuppression. COVID-19 severity was determined in the hospitalized individuals as per the criteria defined by the National Institutes of Health [42], i.e., approximated with respect to the maximum oxygen needed, and the highest level of care provided. Accordingly, the cases were classified as having *mild* (pandemic department, no oxygen supplementation), *moderate* (pandemic department, oxygen supplementation <5L/min), *severe* (pandemic department or intermediate care unit, oxygen need ≥5L/min supplemented by HFNO or CPAP) and *critical* (intensive care unit, with or without mechanical ventilation) COVID-19 illness.

### Sera and Peripheral Blood Mononuclear Cells

Sera were prepared from venous whole blood collected in Vacuette^®^ tubes (Freiner Bio-one GnbH, Kremsmunster, Austria) by centrifugation at 1000 g for 10 minutes at room temperature and stored at -80°C until further use. Peripheral blood mononuclear cells (PBMCs) were isolated from the whole blood by Ficoll-Paque (GE Healthcare, ThermoFisher) density gradient centrifugation. PBMCs were aliquoted in freezing medium (8% DMSO in FBS) and cryopreserved at -150°C for later use in the experiments.

### Flow Cytometry of Whole Blood

The levels of CD3^+^ CD4^+^ T cells and CD3^+^ CD8^+^ T cells/µl were measured using BD Trucount™ Tubes (BD Multitest™ 340491, BD Biosciences) with FITC-conjugated anti-CD3, PE-conjugated anti-CD8, PerCP-conjugated anti-CD45 and APC-conjugated anti-CD4 antibodies (BD Multitest™, 342417, BD Biosciences). Details of additional antibodies used for analysis of T- and B-cell subpopulations are provided in **Supplementary Table 1**. Red blood cells in the whole blood samples were lysed using a commercial BD FACS^TM^ Lysing Solution (349202, BD Biosciences). The samples were run/acquired on a FACScanto II (BD Biosciences). Gating strategies employed for flow cytometry investigations to analyze the cellular clinical parameters are illustrated in **Supplementary Figure 1**.

### Spectral Flow Cytometry

Cryopreserved PBMCs were thawed, washed by centrifugation (1800 RPM, 6 min at 4^°^C), resuspended in FACS buffer (PBS with 0.2%FCS) and counted. Cell density was adjusted to 1×10^6^ cells/ml and added to the FACS tubes (Falcon^®^ Brand, VWR). After pelleting, cells were incubated with FcR blocking reagent (1/15 dilution, Miltenyi), ViaKrome 808 viability dye (1/50 dilution, Beckman Coulter) and CellBlox blocking buffer (1:30 dilution, ThermoFisher), for 20 mins at 4°C in the dark. After washing, the PBMCs were incubated with 30μl of antibody cocktail (See **Supplementary Table 2** for antibodies and conjugation) for 30 min at 4°C. The PBMCs were washed again and resuspended in 200μl FACS buffer and run/acquired on a Cytek^®^ Aurora (Cytek Biosciences, France) flow cytometry system. The spectral flow data was analyzed/processed using OMIQ (OMIQ, Inc, Santa Clara, CA, USA) and FlowJo™ v10.8 Software (BD Life Sciences). Gating strategies used for phenotyping the cells by spectral flow cytometry are illustrated in **Supplementary Figure 2**.

### IFNγ-ELISPOT Assay

96-well plates (Millipore Multiscreen Filtration, Merck Millipore, Sweden) were pre-treated with coating buffer (0.1M Na-Carbonate–Bicarbonate buffer pH 9.5), after which, wells were coated with human anti-IFN-γ monoclonal antibodies (1-D1K, Mabtech Sweden) at a concentration of 5μg/ml and diluted with the same coating buffer. After overnight incubation at 4°C, the wells were washed 4 times with PBS (Cytiva HyClone™ Dulbecco’s PBS, Fisher Scientific), and quenched in 5% PHS in RPMI supplemented with gentamicin (20 µg/ml) and HEPES (10 mM), before seeding the wells with PBMCs at up to 300,000 per well. Later, different stimulants were added, and the plate was incubated for 48 hours at 37°C. The stimulates were pools of 2 µg/ml SARS-CoV-2 peptides (15-mer sequences with 11 amino acid overlaps) consisting of the surface glycoprotein spike (S) (PepTivator^®^ SARS-CoV-2 Prot_S, Miltenyi Biotech); containing the immunodominant sequence domains of the spike, spike plus (S+) (PepTivator^®^ SARS-CoV-2 Prot_S+, Miltenyi Biotech); containing the sequence of a portion of the spike region, or nucleocapsid protein (N) (PepTivator^®^ SARS-CoV-2 Prot_N), Miltenyi Biotech); covering the full protein sequence. Control conditions included mock i.e., PBMCs without stimuli (for background levels) or stimulation with 5 µg/ml phytohemagglutinin (PHA)-M (Sigma-Aldrich) and 20 µg/ml tetanus toxoid (Creative^TM^ Biolabs) (as positive controls).

After 48 hours of incubation, the plates were washed four times with 0.05% Tween 20 in PBS, followed by 2 hours of incubation at 37°C with biotinylated anti-human IFN-γ monoclonal antibodies (clone 7-B6-1, Mabtech) diluted at 1μg/ml in PBS. Subsequently, the plates were washed four times with 0.05% Tween 20 in PBS, and then were incubated with 50μl/well Avidin-Peroxidase-Complex (Vectastain ABC kit: Vector Laboratories, USA) at 37°C for an hour. The Avidin-Peroxidase Complex was prepared and left to stand for 30 min at room temperature before use. The wells were washed 4 times with wash buffer, after which 50μl stable DAB (Invitrogen, Fisher Scientific, Sweden) was added to each well for 3-5 min. Finally, the plates were washed with tap water three times and left to air dry. Spots were counted manually using an inverted phase-contrast microscope (Nikon SMZ1500, Light source Zeiss KL1500 LCD).

### Anti-SARS-CoV-2 Spike and Nucleocapsid IgG

Qualitative measurement of anti-SARS-CoV-2 spike IgG, including anti-spike neutralizing IgG antibodies directed against the receptor binding domain (RBD) of S1 subunit was performed using a commercial chemiluminescent microparticle-based immune assay (Abbott SARS-CoV-2 IgG II Quant/6S60 ARCHITECT SARS-CoV-2 IgG kit) and anti-SARS-CoV-2 nucleocapsid IgG (6R86 ARCHITECT SARS-CoV-2 IgG) using a commercial ARCHITECT Abbott (Abbott Laboratories Diagnostics Division, Abbott Scandinavia AB) kit. The anti-SARS-CoV-2 spike IgG are expressed as standardized binding antibody units (BAU)/mL, calibrated to the WHO International Standards for anti-SARS-CoV-2 immunoglobulin (human) (NIBSC Code 20-136) [43] with a positivity cut-off of 7.1 BAU/mL. The anti-SARS-CoV-2 spike IgG positivity cut-off was 1.4.

### Statistical Analysis

Data and statistical analyses were carried out using GraphPad PRISM v9.0 (GraphPad Software, CA, USA). Differences among the study groups were analyzed with either unpaired, parametric T test with Welch’s correction, or Brown-Forsythe and Welch ANOVA tests, with no correction for multiple comparisons. All differences with P values of <0.05 were considered statistically significant.

## Results

### Clinical Characteristics of COVID-19 Cohort Patients

During a 15 month period (July 2020 to October 2021), blood samples were collected from hospitalized COVID-19 patients and healthy participants. From this cohort, we analyzed data from 46 COVID-19 patients and 31 healthy controls. Furthermore, a subset of donors was followed longitudinally up to 6-7 months. Our analysis focused on the effect of COVID-19 on various T cell subsets and SARS-CoV-2-specific T cell and B cell immune responses (**Figure 1**).

Our hospitalized COVID-19 cohort represents the infection in wider society in terms of biological sex distribution, encompassing two-thirds of males and one third of females [44], and a median age of 56.6 (32 to 91) years (**Table 1**). The disease scores for the patients were moderate or severe as described by the National Institutes of Health [42]. Our control cohort included a majority of females with a median age of 45 years and an age range of 26-62 years. The level of C-reactive protein (CRP) was significantly elevated (88.4 mg/L) in the hospitalized cohort at inclusion with an 8-fold increase as compared to the healthy controls. In addition, the level of lactate dehydrogenase (LDH) was much higher in most patients than in the healthy controls with median of 6.9 µKat/L (**Table 1 and** **Figure 2A**). Both these findings are consistent with observations made by others [45, 46]. No association was found between disease severity and biological sex, age, or clinical characteristics (data not shown).

**Figure 2.**
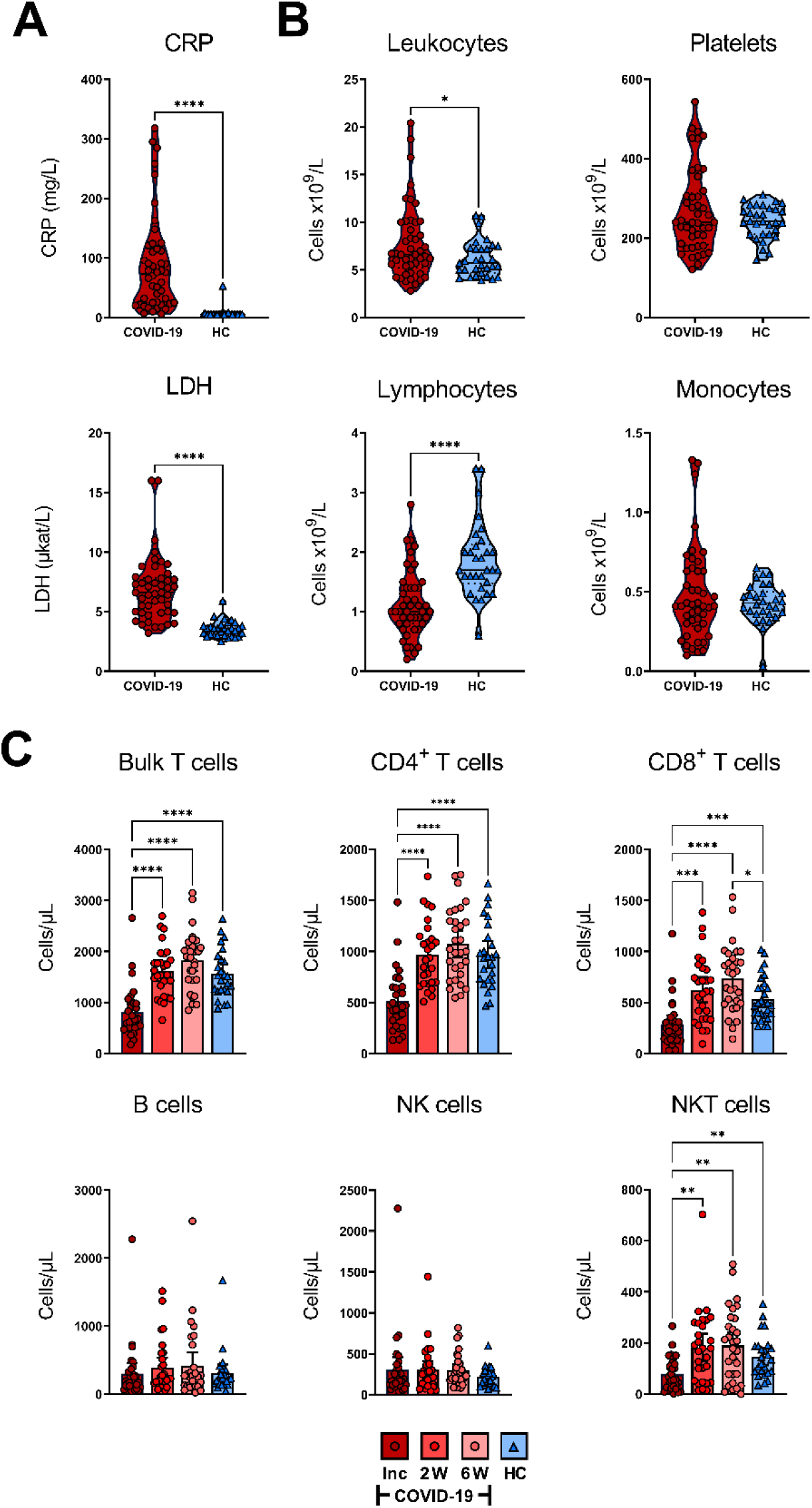
Effects of COVID-19 on clinical parameters and levels of white blood cells and platelets. Peripheral blood was collected from patients with COVID-19 that required hospitalization (N=46) and healthy controls (N=31) and levels of clinical markers and white blood cells were measured at inclusion. (**A**) The levels of CRP (mg/L), and LDH (µKat/L) were measured in blood of COVID-19 patients and healthy controls. (**B**) Leukocytes, lymphocyte, monocyte, and platelet counts in COVID-19 patients and healthy individuals at study inclusion was measured in blood measured as cells/particles x 10^9^/L. (**C**) The absolute number of T cells, CD4^+^ T cells, CD8^+^ T cells, B cells, NK cells, and NKT cells per µL whole blood from COVID-19 patients and healthy individuals were assessed by flow cytometry. The statistical testing was done by unpaired, parametric t test with Welch’s correction, or Brown-Forsythe and Welch ANOVA tests. *p ≤ 0.05, **p ≤ 0.01, ***p ≤ 0.001, ****p ≤ 0.0001. Inc = Inclusion in study at the hospital, 2W = 2 weeks, 6W = 6 weeks, HC = healthy control.

### Effect of COVID-19 on the Levels of Circulating White Blood Cells and Thrombocytes

The number of lymphocytes/L was significantly decreased, and leukocytes/L was significantly increased at inclusion compared to the healthy controls, whereas monocyte and platelet counts were not significantly affected, although there was a minor fraction of COVID-19 patients with increased levels of platelets (**Figure 2B**). To further characterize the attrition in lymphocytes, we analyzed CD4^+^ T cells, CD8^+^ T cells, B cells, NK cells, and NKT cells in peripheral blood. There was a significant decrease in T cells, CD4^+^ T cells, CD8^+^ T cells, and NKT cells/µL at inclusion among COVID-19 patients relative to healthy controls. Furthermore, the lymphocyte populations experienced recovery with a significant increase in cell numbers at week 2, which remained stable until the 6 week time point (**Figure 2B**). Regarding the CD8^+^ T cells, the increase was significantly higher at week 6 compared to healthy controls. We found no significant alteration in the absolute number of NK cells or B cells (**Figure 2B**). Together, these data are reflective of the alterations in the levels of various immune cell types during COVID-19, in line with observations made by others [25, 47–50].

### Hospitalized SARS-CoV-2-Infected Individuals have Prolonged Reduction of CD3^+^ T cells but CD8^+^ T cells Gradually Rebound from 6 Weeks Onwards

To assess the longitudinal effect of COVID-19 on T cell populations, we sampled PBMCs from 21 patients with COVID-19 and 16 healthy controls over a period of 6-7months. PBMCs were analyzed by flow cytometry to investigate immunophenotype across the various T cell compartments. The percentages of CD3^+^ T cells were significantly decreased in patients at inclusion compared to the later time points and to healthy controls (**Supplementary Figure 3A**). At week 2, the fraction of CD3^+^ T cells was still decreased in comparison with the healthy controls. While comparable percentages of CD4^+^ T cells were observed at hospitalization and week 2, there was a significant decrease in the levels after 6 weeks and 6-7 months, relative to the healthy controls (**Supplementary Figure 3B**). Conversely, CD8^+^ T cells were significantly increased at 6 weeks and 6–7 months, in comparison to healthy individuals, while similar levels of these cells were seen at the first two time points (hospitalization and 2 weeks) (**Supplementary Figure 3C**).

### COVID-19 Patients have a Sustained, Elevated CD8^+^ TEMRA Population

The environment locally and systemically created by an infection can give rise to alterations in the general memory T cell populations [51]. Here, we examined the effect COVID-19 had on naïve and memory CD4^+^ and CD8^+^ T cells (**Figure 3**). Distinct differences in the CD4^+^ and CD8^+^ T populations between a COVID-19 patient and a healthy control are shown in the representative pseudo-color dot plots (**Figure 3A**). A slight, but not significant, decrease was seen in the naïve fraction of the CD4^+^ TH cells over time, with no alterations in the CD4+ TEMRA (CCR7^-^CD45RA^+^) or T cell effector memory (TEM; CCR7^-^CD45RA^-^) fractions (**Figure 3B**). There was, notwithstanding, a significant upregulation in the T cell central memory (TCM; CCR7^+^CD45RA^-^) compartment of CD4^+^ T cells at 6-7 months in comparison with hospitalization. We found that CD8^+^ T cells displayed more sustained alterations in their naïve and TEMRA compartments compared with healthy controls. However, the frequencies of TCM and TEM were not affected (**Figure 3B**). A gradual decrease over time occurred in the naïve CD8^+^ T cells of COVID-19 patients, with the 2 weeks, 6 weeks, and 6–7 month time points presenting significantly decreased fractions of these cells compared to healthy controls (**Figure 3B**). In contrast, we found a sustained increased percentage of CD8^+^ TEMRA cells throughout the investigation, in comparison to healthy controls.

**Figure 3.**
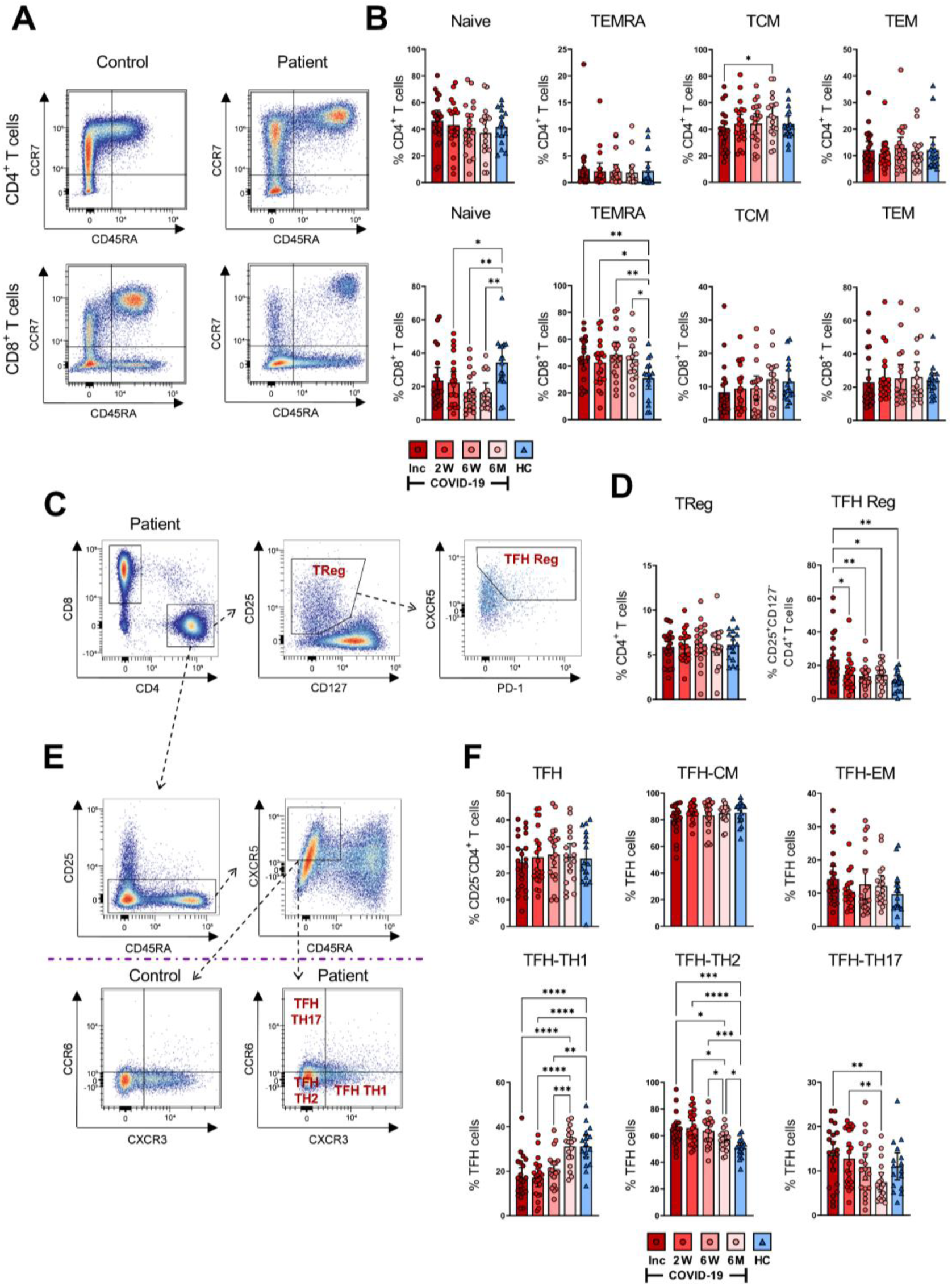
SARS-CoV-2-specific CD4^+^ and CD8^+^ T cell subsets during COVID-19 infection. PBMCs were collected from COVID-19 patients that needed hospitalization (N=23) and healthy controls (N=16) over a 6-7 month period. Cells were stained with antibodies for flow cytometry to immunophenotype the various T cell populations. (**A**) Representative flow plot of CD4^+^ (upper panel) and CD8^+^ (lower panel) naïve and memory T cell populations. (**B**) Percentages of naïve (CCR7^+^CD45RA^+^), TEMRA (CCR7^-^CD45RA^+^), TCM (CCR7^+^CD45RA^-^), and TEM (CCR7^-^CD45RA^-^) subsets in CD4^+^ (upper panel) and CD8^+^ (lower panel) T cells. (**C**) Representative gating strategy of CD4^+^ Tregs (CD25^+^CD127^-^) and TFH Reg (CXCR5^+^PD-1^+^CD25^+^CD127^-^) on CD4^+^ T helper cells. (**D**) Fractions of Tregs (left panel) and TFH Tregs (right panel). (**E**) Representative gating and (**F**) proportions of TH1, TH2, and TH17 TFH T cells. Data is represented as mean with 95% Cl, with significance of *p≤0.05, **p≤0.01, ***p≤0.001, ****p≤0.0001., determined using Brown-Forsythe and Welch ANOVA tests. Inc = Inclusion in study at the hospital, 2W = 2 weeks, 6W = 6 weeks, 6M = 6-7 months, HC = healthy control.

Next, we characterized the regulatory T cell (Treg; CD25^+^CD127^-^) fraction of CD4^+^ T cells (**Figure 3C** and **3D**) and found comparable levels of CD4^+^ Tregs in patient and control samples (**Figure 3D**). The TFH subset amongst Tregs was significantly increased in COVID-19 patients at hospitalization but decreased to a comparable level relative to healthy controls by the 2 week time point (**Figure 3D**). Overall, we found alterations in the memory T cell pool and this was most evident in the CD8^+^ TEMRA subset, where the changes were sustained until the end of the study.

### COVID-19 Gives Rise to Altered TFH Cell Subsets with Consistent and Elevated TH2 Phenotype

Follicular helper T cells interact with B cells and facilitate the humoral arm of immunity by enhancing B cell responses, ultimately resulting in the generation of long-lasting, high-affinity antibodies, especially in the context of viral infections [52–54]. We evaluated TFH responses (**Figure 3E** and **3F**) and found that TCM and TEM levels among the TFH remained unchanged (**Figure 3F**). In contrast, the TFH subsets had significant alterations in the TH1 (CXCR3^+^CCR6^-^), TH2 (CXCR3^-^CCR6^-^) and TH17 (CXCR3^-^CCR6^+^) proportions (**Figure 3F**). There was a significant drop in CD4^+^ TFH-TH1 cells at hospitalization among COVID-19 patients as compared to healthy controls, which consistently increased throughout the infection, reaching a level at 6-7 months that was comparable to healthy controls (**Figure 3F**). Conversely, TFH-TH2 cells were significantly upregulated at hospitalization in comparison to healthy controls. The fractions of these cells were consistently elevated during the infection. Although their levels eventually decreased, they remained higher compared to healthy controls after 6-7 months (**Figure 3F**). The level of TFH-TH17 cells was not significantly different from the control group at any time point, despite appearing lower after 6-7 months. Nonetheless, there was a gradual decrease of TFH-TH17 over time, which reached significance by the final time point (**Figure 3F**). There was a clear change in the TFH population, with a sustained increase in the TH2 subset in particular.

### COVID-19 Patients have Activated T Cells at Inclusion, and Certain Cell Populations such as CD69^+^ CD4^+^ T Cells are Maintained Throughout the Study

Classical markers of early and late activation of T cells are represented by CD69 and CD38/HLA-DR, respectively [37, 38]. Persistent expression of these markers is suggestive of a hyper-immune activation state and affirms an ongoing phase of systemic inflammation and tissue injury occurring in severe COVID-19 [55]. Hence, to examine the activation status of CD4^+^ and CD8^+^ T cells, we assessed the cells for CD38^+^HLA-DR^+^ and CD69^+^ expression (**Figure 4**). We found clear differences in the activation status of the T cells between the COVID-19 patient and healthy control as illustrated in the representative scatter plots (**Figure 4A**). The fraction of CD38^+^HLA-DR^+^ activated CD4^+^ T cells in the COVID-19 patients significantly decreased over the study duration and appeared to be lower than healthy controls at 6-7 months although this was not significant (**Figure 4B**). The levels of CD69^+^ activated CD4^+^ T cells were significantly higher at hospitalization than in healthy controls. This elevation was sustained until the 6-7 month time point despite having dropped significantly from the initial levels (**Figure 4B**).

**Figure 4.**
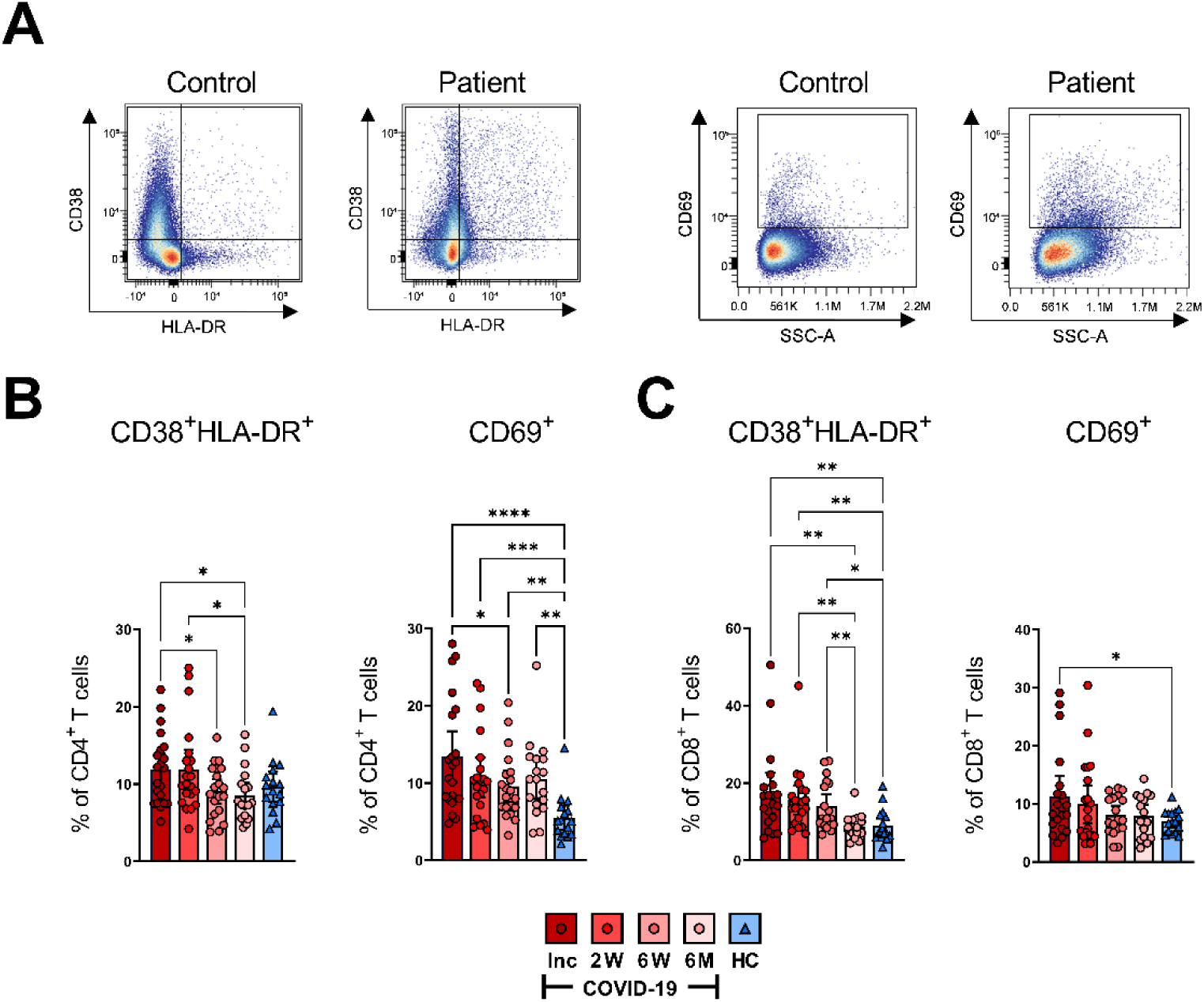
T cells are activated during SARS-CoV-2 infection. PBMCs collected from COVID-19 patients that required hospitalization (N=23) and healthy donors (N=16), were stained for flow cytometry to assess the activation status of CD4^+^ and CD8^+^ T cells. (**A**) Representative dot plots for CD38^+^/HLA-DR^+^ and CD69^+^ activated CD4^+^ and CD8^+^ T cells. (**B**) Proportions of the CD38^+^/HLA-DR^+^ and CD69^+^ activated CD4^+^ T cells. (**C**) Proportions of the CD38^+^/HLA-DR^+^ and CD69 activated CD8^+^ T cells. Data is represented as mean with 95% Cl, with significance of *p≤0.05, **p≤0.01, ***p≤0.001, ****p≤0.0001, determined using Brown-Forsythe and Welch ANOVA tests. Inc = Inclusion in study at the hospital, 2W = 2 weeks, 6W = 6 weeks, 6M = 6-7 months, HC = healthy control.

Activated CD8^+^ T cells in COVID-19 patients were also abundantly increased at hospitalization when compared to healthy controls. We observed a decrease over the 6-7 months period to the same level as healthy controls, although the CD38^+^HLA-DR^+^ phenotype was maintained up to 6 weeks where the CD69^+^ phenotype declined earlier (**Figure 4C**). The data collectively illustrates a significantly increased activation of both CD4^+^ and CD8^+^ T cells in COVID-19 patients at the initial sampling period that is sustained for some but not all markers in each subset.

### COVID-19 Induces a Long Term Altered Exhausted CD57^+^CD8^+^ T Cell Population

To investigate the magnitude of T cell impairment during SARS-CoV-2 infection and recovery, we measured the frequency of co-inhibitory markers that are normally upregulated during exhaustion and suppression on CD4^+^ and CD8^+^ T cell subsets. CD57 is a marker of terminally differentiated non-proliferative T cells (senescence and exhaustion) and the frequency of CD57^+^CD4^+^ and CD57^+^CD8^+^ T cells increases with age as well as with cancer and chronic infections [56]. Both CD4^+^ and CD8^+^ T cell populations were examined for expression of CD57 as shown by a representative pseudo color dot plot (**Figure 5A**). We found no significant difference in the proportion of CD57^+^CD4^+^ T cells between COVID-19 patients and healthy controls (**Figure 5B**). Interestingly, we observed a significant surge in the percentage of CD57^+^CD8^+^ T cells among COVID-19 patients at inclusion compared to healthy controls, which increased further by the 6 week time point and was sustained up until 6-7 months (**Figure 5B**). Taken together, our data clearly shows a long-lasting effect on the CD8^+^ T cell population with increased level of exhausted CD57^+^ CD8^+^ T cells.

**Figure 5.**
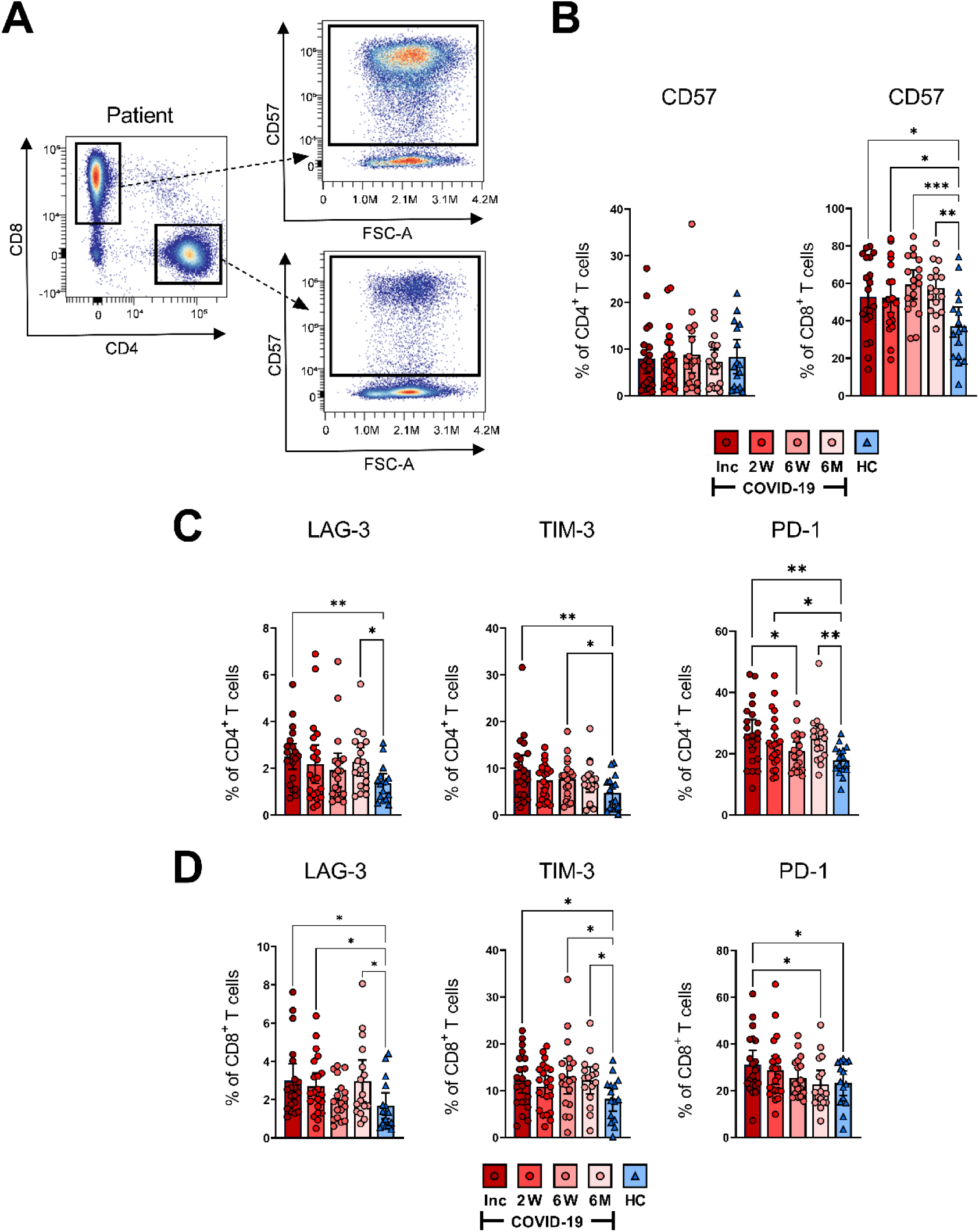
T cell impairment associated with COVID-19. PBMCs collected from COVID-19 patients that needed hospitalization (N=23) and healthy controls (N=16), were stained for flow cytometry to assess the expression of negative immune checkpoint factors and exhausted status of CD4^+^ and CD8^+^ T cells. (**A**) Representative flow cytometry plots show how CD57 expression levels were determined for CD4^+^ and CD8^+^ T cells. (**B**) Expression frequency of CD57 among CD4^+^ and CD8^+^ T cells. (**C**) Expression frequencies of LAG-3^+^, TIM-3^+^ and PD-1^+^ among CD4^+^ and (**D**) CD8^+^ T cells. Data is represented as mean with 95% Cl, with significance of *p≤0.05, **p≤0.01, ***p≤0.001, determined using Brown-Forsythe and Welch ANOVA tests. Inc = Inclusion in study at the hospital, 2W = 2 weeks, 6W = 6 weeks, 6M = 6-7 months, HC = healthy control.

### COVID-19 Gives Rise to High Expression of LAG-3, TIM-3 and PD-1 that is Maintained for a Prolonged Duration, Indicative of Suboptimal T Cell Responsiveness

Next, we examined the expression of three co-inhibitory markers i.e., LAG-3, which inhibits T cell activation and promotes a suppressive phenotype [57]; TIM-3, a T cell exhaustion marker [58] and PD-1, which down-regulates functional T cell activation [59]. These markers of T cell impairment are all expressed during chronic infections. We observed a significant increased level of LAG-3^+^CD4^+^ T cells among COVID-19 patients at hospitalization and again at the 6-7 month time point when compared to healthy controls (**Figure 5C**). Similarly, the proportion of TIM-3^+^CD4^+^ T cells was elevated at hospitalization, and again at the 6 week time point. At the 6-7 month time point, the level appeared to rise but was not significant (**Figure 5C**). The percentage of PD-1^+^CD4^+^ T cells was significantly raised compared to healthy controls at all time points except for a significant dip at 6 weeks (**Figure 5C**).

A significant elevation of LAG-3^+^CD8^+^ T cells was observed at hospitalization and again after 2 weeks, and 6-7 months (**Figure 5D**). The fraction of TIM-3^+^ CD8^+^ T cells was significantly raised at hospitalization, 6 weeks, and 6-7 months (**Figure 5D**). The levels of PD-1^+^ CD8^+^ cells were significantly raised at hospitalization and reverted to normal levels over the following 6-7 months (**Figure 5D**). Overall, there was a general trend within all three markers of being elevated among COVID-19 patients at hospitalization when compared to healthy controls across both CD4^+^ and the CD8^+^ T cell subsets. Of note, there was a pattern within most of the data, of levels decreasing after the first time point, but increasing again by the 6–7 month time point.

### COVID-19 Patients Develop Spike and Nucleocapsid SARS-CoV-2-Specific T Cells with the Ability to Respond with IFN-γ Production Following Stimulation

ELISPOT assay measures the antigen-specific responses of T cells [60–62], and has been used to determine the functional responses of SARS-CoV-2-specific T cells [63–65]. Here, we evaluated the level of SARS-CoV-2-specific IFN-γ producing bulk T cells in the COVID-19 patients, following stimulation with overlapping peptides covering the spike (containing the immunodominant RBD), spike+ (containing a portion of the spike region) and the nucleocapsid (covering full sequence) proteins (**Figure 6**). We found SARS-CoV-2 specific T cell responses in all COVID-19 patients (**Figure 6** and **Supplementary Figure 4**). The SARS-CoV-2 specific response from inclusion to the 6–7 month time point in a representative donor is presented in **Figure 6A**. We observed a gradual and significant increase in the number of spike (**Figure 6B**), spike+ (**Figure 6C**), and nucleocapsid (**Figure 6D**) SARS-CoV-2-specific T cells, which was sustained for 6-7 months. The antigen specific T cell responses in individual donors are illustrated in **Supplementary Figure 4**. Altogether, we found the frequency of bulk SARS-CoV-2-specific T cells to be increased over the duration of the investigation.

**Figure 6.**
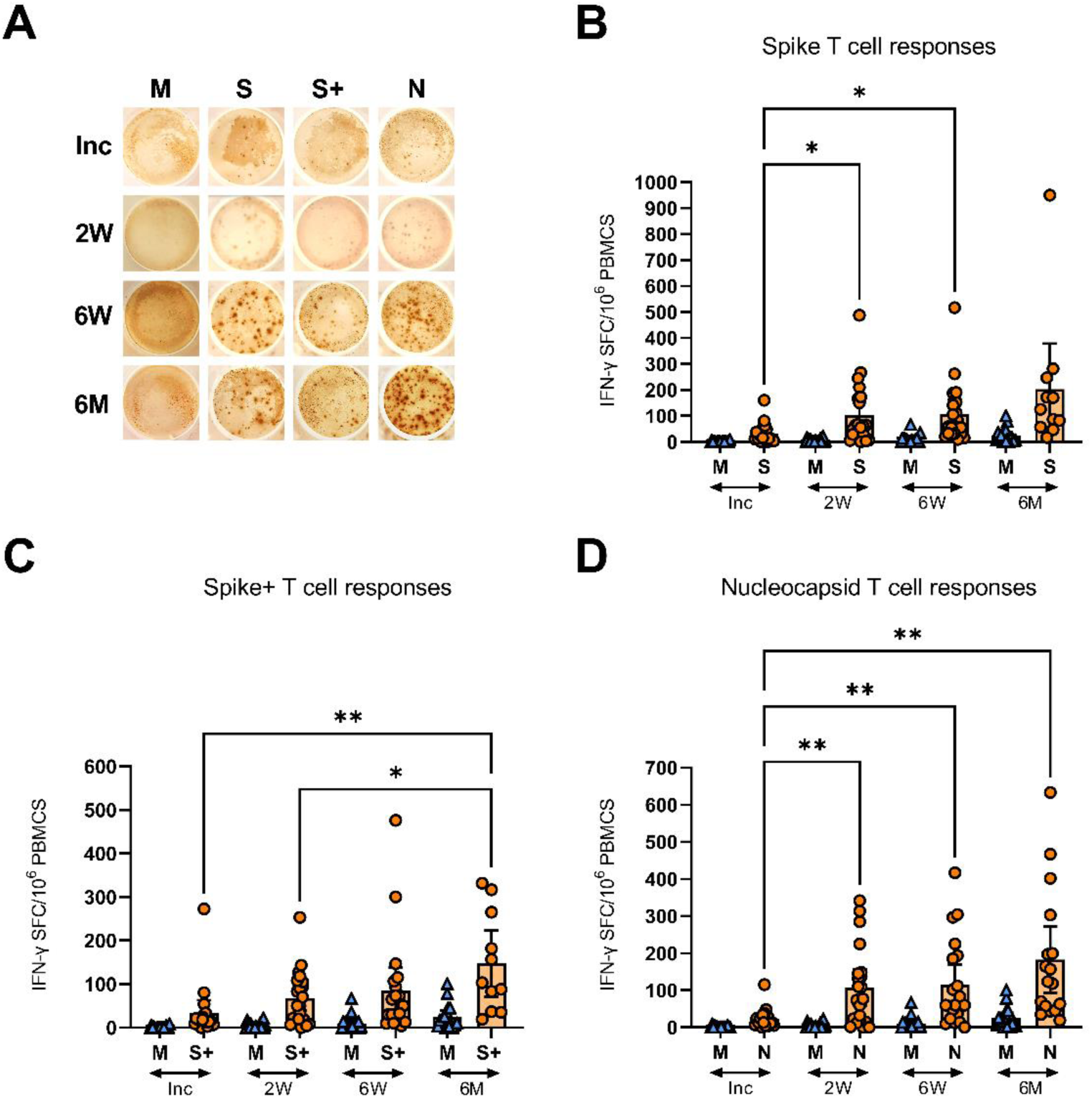
SARS-CoV-2-specific IFN-γ T cell responses in COVID-19 patients. PBMCs collected from COVID-19 patients that needed hospitalization (N=23) were left unstimulated (M) or incubated with SARS-CoV-2 specific peptides: Spike (S), spike+ (S+), and nucleocapsid (N); and assessed for their production of IFN-γ by ELISPOT. Representative images of the IFN-γ production in a COVID-19 hospitalized patient at inclusion, 2 weeks, 6 weeks, and 6-7 months, each dot represent one antigen-specific T cell (**A**). Antigen specific T cells responses as spot forming cells (SFC) per 10^6^ PBMCs, specific for spike (**B**), spike+ (**C**), and nucleocapsid antigens (**D**). Data is represented as mean with 95% Cl, with significance of *p ≤ 0.05, **p ≤ 0.01, determined using Brown-Forsythe and Welch ANOVA tests. Inc = Inclusion in study at the hospital, 2W = 2 weeks, 6W = 6 weeks, 6M = 6-7 months, HC = healthy control.

### COVID-19 Patients Develop SARS-CoV-2-Specific Antibody Responses and have Activated B Cells Early in the Course of the Disease

The levels of anti-SARS-CoV-2 nucleocapsid and spike-neutralizing Abs in the COVID-19 hospitalized patients were assessed at inclusion, 2 weeks, 6 weeks, and 6-8 months. The levels of antibodies increased significantly from inclusion to the 2 week time point and were also significantly higher at 6 weeks, before declining back to inclusion levels by 6-8 months (**Figure 7A**, left). The spike-nAbs were of the same profile as the anti-nucleocapsid Abs, which increased significantly from inclusion compared to the 2 and 6 week time points and eventually declined at the 6-8 month time point (**Figure 7A**, right). We ensured that the SARS-CoV-2 vaccinated participants were not tested at the final time point.

**Figure 7.**
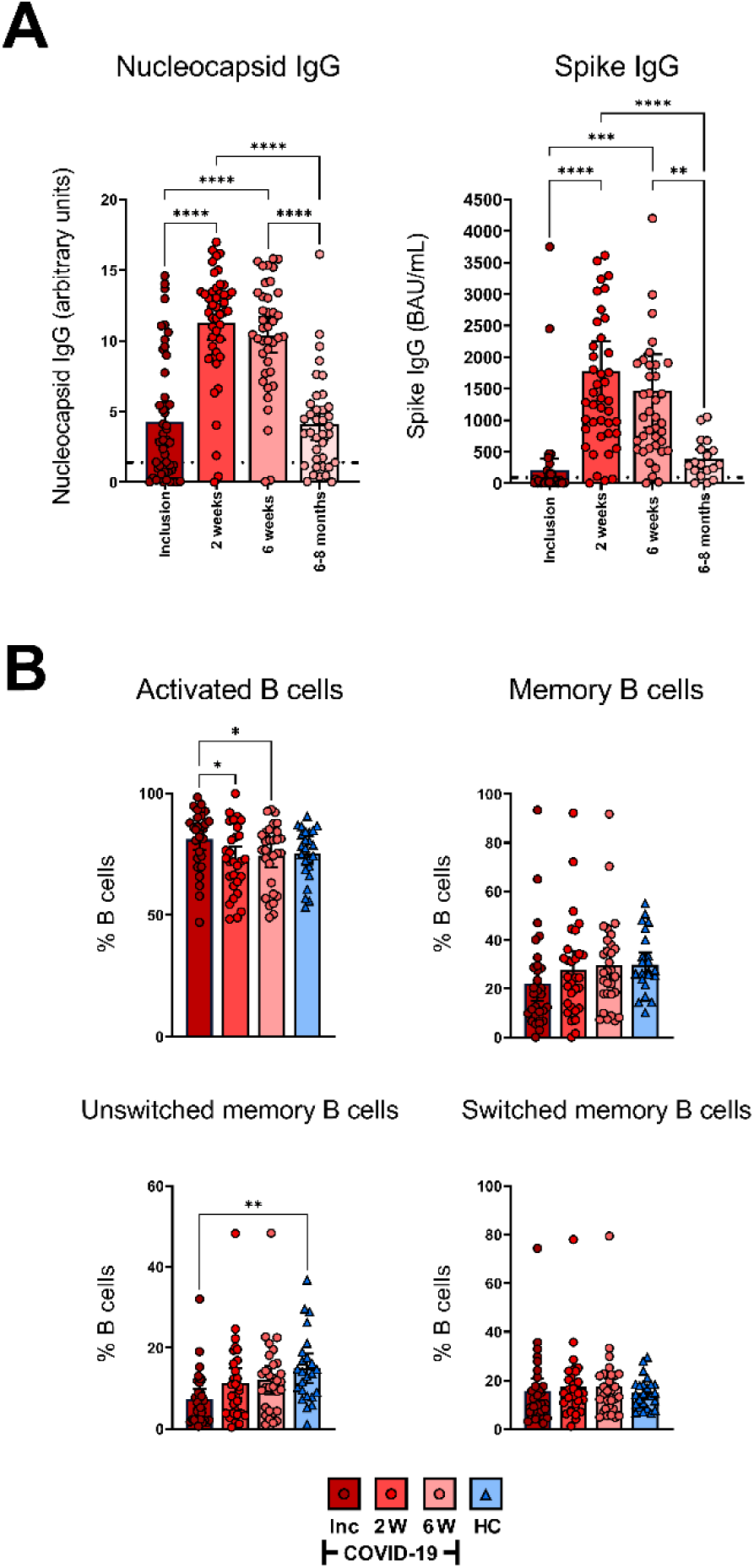
All COVID-19 patients develop SARS-CoV2 specific antibody responses and have activated B cells early in the disease. Sera were collected from COVID-19 patients that required hospitalization (N=46) and healthy donors (N=31). (**A**)The levels of nucleocapsid antibodies (Units, 0.8-1.4 borderline positive and > 1.4 for positive result) (left) and neutralizing spike antibodies (BAU/ml, ≥ 30 for positive result) (right) were measured in sera from inclusion up to 6-8 months. (**B**) The B cell populations were measured by flow cytometry as percentage out of the CD19^+^ B cells; CD19^+^CD27^+^ B cells (memory B cells), CD19^+^CD27^+^IgD^+^ B cells (unswitched B cells), CD19^+^CD27^+^IgD^-^ B cells (Switched B cells), and CD19^+^CD38^+^ B cells (activated B cells). Data is represented as mean with 95% Cl, with significance of *p ≤ 0.05, **p ≤ 0.01, ***p ≤ 0.001, ****p ≤ 0.0001, determined using Brown-Forsythe and Welch ANOVA tests. Inc = Inclusion in study at the hospital, 2W = 2 weeks, 6W = 6 weeks, HC = healthy control.

The frequency of activated B cells, defined by CD38 expression [66, 67], was significantly higher at inclusion compared to the 2 and 6 week time points (**Figure 7B**). The levels of memory B cells at inclusion were reduced, but not significantly so, when compared to healthy controls. Similarly, unswitched memory B cell levels were significantly decreased at hospitalization. The percentages of both the populations increased at the 2 week time point to levels comparable to healthy controls (**Figure 7B**). There were no significant alterations observed in the levels of switched memory B cells (**Figure 7B**). Given there was no effect on the absolute numbers of B cells per µl (**Figure 1C**), these data indicate an alteration in the proportion of the different sub-populations within the B cell compartment. Collectively, the data illustrate a decrease in the unswitched B cell population and subsequent waning of SARS-CoV-2-specific antibodies over time.

## DISCUSSION

We have profiled T cell populations during COVID-19 in a longitudinal study cohort, where we followed them from inclusion up to 6-8 months post-recovery, using spectral flow cytometry and serological and immunological assays. Additionally, we have evaluated the SARS-CoV-2 specific T cell responses, B cell responses, and SARS-CoV-2 antibodies generated in the cohort. We found wide-ranging alterations to the T cell compartment including a rise in CD8^+^ TEMRA and TFH TH2 cells, and different forms of functional impairments that lasted long into convalescence.

Our initial measurement of clinical parameters was consistent with others showing elevated CRP and LDH, common acute-inflammatory biomarkers [49, 68, 69]. These have been shown to have a positive correlation with the development of pulmonary lesions during early COVID-19, and consequently with severe COVID-19 and related respiratory complications [70, 71]. We recently found an association of soluble urokinase Plasminogen Activator Receptor with COVID-19 severity, and length of hospitalization [72]. Additionally, lymphopenia was observed in many of the COVID-19 patients in our cohort at inclusion (i.e., at hospitalization), and is a common clinical observation [73–76] that may likely be attributed to the cells relocating or consequently dying at this stage of the disease. Indeed, a highly inflammatory form of cell death, i.e., pyroptosis, induced in infected and uninfected cells, appears to be a major contributing factor for the onset of strong inflammatory responses seen globally in many individuals with COVID-19 [8, 49, 69, 77, 78]. Furthermore, we and others have shown that the T cell compartment is highly affected, and to a higher degree than immune cells such as B or NK cells [79–82], which implies that T cell subsets play a paramount role in COVID-19 pathogenesis.

Memory T cells, both general and antigen-specific, in the context of SARS-CoV-2 infection have been widely studied, notwithstanding often for shorter time-periods [22, 75, 83, 84] with fewer long-term studies, i.e. 8-9 months [85]. Evidence from the SARS-CoV outbreak in 2005 suggests that anti-SARS-CoV antibodies fell below detection limits within two years [86], and SARS-CoV-specific memory T cells were detectable 11 years after the SARS outbreak [87]. Memory T cells are an important and diverse subset of antigen experienced T cells that are sustained long-term, and when needed are converted into effector cells during re-infection/exposure [88, 89]. Depending on their cellular programming and phenotype they are classified into different central and effector memory subtypes. The effector subsets contain the CD45RA^+^CCR7^-^ TEMRA, which are essentially TEM that re-express CD45RA after antigen stimulation [90]. Not much is known about the functionality of this population, but CD4^+^ TEMRA are implicated in protective immunity [90]. Furthermore, high levels of virus specific CD8^+^ TEMRA are maintained after dengue vaccination [91]. We found an elevation of CD8^+^TEMRA that lasted throughout the study, i.e., 6-7 months, which contrasted with the CD4^+^ TEMRA that remained unaltered by COVID-19. Previous studies have shown the CD8^+^ TEMRA population to be increased at hospitalization [92, 93] and sustained for 6 weeks [92]. Nonetheless, as far as we are aware, no studies have followed up beyond the 6 week time point. Currently, the exact role of CD8^+^ TEMRA in COVID-19 remains largely ambiguous, but Cohen et al [85] found an increase in SARS-CoV-2 specific CD8^+^ TEMRA over time. In our case, we have explored the whole expanded CD8^+^ TEMRA population and cannot confirm if there was a larger fraction of antigen-exposed, i.e., SARS-CoV-2-specific T cells, among the population.

We found that all COVID-19 patients developed SARS-CoV-2 antigen specific T cells, that increased over time. Our findings are in accordance with other studies that have shown that SARS-CoV-2 infection results in increased expansion of antigen specific CD4^+^ and CD8^+^ T cell subsets [63, 94]. It is still unclear if the lower antigen specific responses seen at 2 and 6 weeks compared to 6-7 months are due to an overall immunosuppression [95] or a natural development of the immune response over time [65]. Furthermore, there might be a relocation of the cells, as the composition of peripheral antigen specific T cells may not necessarily be indicative of the frequency of the cell phenotypes in the pulmonary compartment in COVID-19.

The TFH cell subsets are essential for inducing high-affinity antibodies [27–31]. The frequency of circulating TFH have been shown to be correlated with neutralizing antibodies in viral infections such as HIV-1 [34], and now SARS-CoV-2 [22, 25, 35]. We found no alteration in the levels of TFH in general, while the frequencies of TFH TH2 were elevated which is consistent with findings by Juno et al [23]. In addition, the TFH regulatory T cells were elevated at inclusion in the study but reverted to normal levels after a few weeks.

Our study clearly shows that activation of CD4^+^ and CD8^+^ T cells is a key finding in COVID-19 patients during active disease, and this is in agreement with others [39, 55]. Concerning the activated CD8^+^ T cell population expressing CD38 and HLA-DR, the levels were elevated compared to healthy controls and were sustained for more than 6 weeks, whereas the CD4^+^ T cells decreased over time within the COVID-19 patients. High levels of HLADR^+^ CD38^+^ CD8^+^ T cells that persisted 30-60 days were detected in individuals with severe COVID-19 [39], which is in accordance with our findings. We found elevated levels of activated CD69^+^ CD4^+^ T cells that were sustained at the 6-7 month follow-up, while the levels of CD69^+^CD8^+^ T cells dropped to healthy control levels already at the 2 and 6 week time points. CD69 expression on T cells is considered an early activation marker [38], and is also implicated in regulating mucosal inflammation [96]. It has also been shown that sustainment of CD69 on circulating CD8^+^ T cells eventually could also accelerate their destruction in the liver [97], and hence it remains to be seen if the activated T cells would still be recruited into the effector or memory pool.

The hyper-activation of T cells is a well-studied mechanism in chronic viral infections [98] and is often dependent on the levels of viral persistence in the circulation. The reason for the prolonged activation of T cells in COVID-19 could be owing to the persistence of viral antigens in the bone marrow, that indirectly affect the circulating lymphocytes [99]. In addition, this may also be attributed to an ongoing chronic inflammation resulting from fibrosis/on-going tissue repair in the lung of patients’ recovering from COVID-19 [100, 101]. Indeed, individuals that had been infected with SARS-CoV experienced dysregulation in lung function that persisted for 2 years [102], and hence in our cohort, 6-7 months is seemingly too short to witness complete recovery. Chronic activation of T cells leads to impairment in their functionality and elevated levels of negative immune checkpoint molecules, which compromise both the antigen specific and bystander responses [103]. The negative immune checkpoint molecules PD-1, LAG-3, or TIM-3 expressing CD4^+^ and CD8^+^ T cells were elevated at inclusion. Additionally, the levels of LAG-3 and PD-1 on CD4^+^ T cells, and LAG-3 and TIM-3 on CD8^+^ T cells were elevated at the 6-7 month time point. The expression of negative immune checkpoint molecules TIM-3 and PD-1 mediates immunosuppression in lung, and was found to be a hallmark of severe COVID-19, particularly in men [104]. Besides the induction of the negative immune checkpoint molecules, chronic stimulation can give rise to CD57^+^ exhausted T cells. In addition, the frequency of CD57^+^ T cells increases with age [56]. There was a high impact on the CD8^+^ T cell population with an increase in CD57^+^CD8^+^ T cells that lasted throughout the study (6-7 months), although we did not find any effect on the CD4^+^ T cell population. Increased CD57 CD8^+^ T cells has previously only been shown early during hospitalization in COVID-19 patients [41].

Today it is clear that all immunocompetent individuals develop SARS-CoV-2-specific antibodies, which is also evident in our investigation. Some studies provide clear evidence that these antibodies are detected only for a few months after infection [105, 106], whereas others support a minimum of 6 months [20, 107]. In our cohort, we found that the SARS-CoV-2 nucleocapsid-specific and spike-specific neutralizing antibody levels lasted 6-8 months post infection, even though there was a drastic decline at some time point after 6 weeks. Our findings are supported by Björkander et al. [94], who have reported that the antibody responses lasted up to 8 months among young adults.

With the continued burden of the current COVID-19 pandemic on the population, there is still a need for more insight into the SARS-CoV-2-specific immune response elicited during infection. Despite having multiple approved and licensed vaccines, the emergence of variants with multiple mutations [108], and the long list of long-term symptoms following a natural SARS-CoV-2 infection [109–111], are still cause for great concern. Further, given the likely impact of inter-human variations in clinical parameters, more data is needed regarding the durability and sustenance of SARS-CoV-2 specific antibodies and T cells generated during COVID-19 and their contribution to the quality of immune responses and what the lasting effects on the immune cells compartment in individual that have recovered from COVID-19.

It is important to note that the current investigation has limitations. A critical factor to consider is the pre-selection of patients for this cohort, with most of the patients included already having moderate to severe COVID-19. It is also noteworthy to consider that health systems vary across geographical locations so patients in other counties might be more diverse due to different thresholds of illness required for hospital admission. Additionally, due to severe disease presentations, there were challenges with obtaining clinical samples from some patients and, in some cases, the cell numbers were low and/or of poor quality. During the study some COVID-19 patients were vaccinated against SARS-CoV-2 and therefore were excluded from our data set as this interfered with some results.

Despite the immensely challenging conditions during the pandemic, we do believe that the cohort presented in this investigation is well characterized and of high quality and value. Altogether, this study highlights the alterations in the immune response incurred during hospital treated SARS-CoV-2 infection and convalescence. These novel longitudinal data illustrate the substantial changes to the T cell landscape, with a persistent increase in markers of activation, exhaustion and senescence lasting for more than 6 months. We further showed an accompanying decay in SARS-CoV-2-specific antibody responses at the same time. Our findings, in combination with others, are valuable in providing insight into SARS-CoV-2 cellular and humoral immunity, and open new avenues to be explored for improved understanding of the long-term alterations described herein in COVID-19 immunopathogenesis.

## Authorship contribution

M.G., F.R.H., C.S., R.G., M.H., and M.L. conducted experiments. M.G., F.R.H., C.S., R.G., M.H., S.N., and M.L. analyzed the data. M.G., F.R.H., E.M.S., S.N., and M.L were involved in the writing of the initial manuscript, and M.G., F.R.H., C.S., R.G., M.H., A.N., J.S., S.N., E.M.S., and M.L. helped in the revision of the manuscript. M.L. designed the experiments. A.N., J.S., S.N., and M.L. were involved in establishing the cohort study. All authors read and approved the final manuscript.

## Acknowledgements

We sincerely thank all the donors who participated in the study. We are grateful to all those who contributed to the study, especially the health care personnel at the Clinic of Infectious Diseases and at the Intensive Care Unit at the Vrinnevi Hospital, Norrköping, Sweden, and the cohort study coordinators. We would also like to thank Annette Gustafsson for study coordination and collecting samples from the donors, and Mario Alberto Cano Fiestas for assisting with processing of samples. We are grateful for the assistance from the Core Facility Flow Unit at Linkoping University, especially Jörgen Adolfsson. We thank Liselotte Ydrenius and Malin Siljhammar and the staff at the Flow Cytometry Unit at Clinical Immunology and Transfusion Medicine, Region Östergötland.

## References

1. WHO. WHO Coronavirus (COVID-19) Dashboard. 2022 1 March [cited 2022 2 March]; Website]. Available from: https://covid19.who.int/.

2. Shang, J., et al., Cell entry mechanisms of SARS-CoV-2. Proc Natl Acad Sci U S A, 2020. 117(21): p. 11727–11734.

3. Hoffmann, M., et al., SARS-CoV-2 Cell Entry Depends on ACE2 and TMPRSS2 and Is Blocked by a Clinically Proven Protease Inhibitor. Cell, 2020. 181(2): p. 271–280 e8.

4. Phua, J., et al., Intensive care management of coronavirus disease 2019 (COVID-19): challenges and recommendations. Lancet Respir Med, 2020. 8(5): p. 506–517.

5. Chen, N., et al., Epidemiological and clinical characteristics of 99 cases of 2019 novel coronavirus pneumonia in Wuhan, China: a descriptive study. Lancet, 2020. 395(10223): p. 507–513.

6. Elezkurtaj, S., et al., Causes of death and comorbidities in hospitalized patients with COVID-19. Scientific Reports, 2021. 11(1): p. 4263.

7. Zhang, B., et al., Clinical characteristics of 82 cases of death from COVID-19. PLOS ONE, 2020. 15(7): p. e0235458.

8. Li, S., et al., SARS-CoV-2 triggers inflammatory responses and cell death through caspase-8 activation. Signal Transduct Target Ther, 2020. 5(1): p. 235.

9. Tay, M.Z., et al., The trinity of COVID-19: immunity, inflammation and intervention. Nat Rev Immunol, 2020. 20(6): p. 363–374.

10. Paolini, A., et al., Cell Death in Coronavirus Infections: Uncovering Its Role during COVID-19. Cells, 2021. 10(7): p. 1585.

11. Alefishat, E., et al., Immune response to SARS-CoV-2 variants: A focus on severity, susceptibility, and preexisting immunity. Journal of Infection and Public Health, 2022. 15(2): p. 277–288.

12. El Zein, S., et al., SARS-CoV-2 infection: Initial viral load (iVL) predicts severity of illness/outcome, and declining trend of iVL in hospitalized patients corresponds with slowing of the pandemic. PLOS ONE, 2021. 16(9): p. e0255981.

13. Rydyznski Moderbacher, C., et al., Antigen-Specific Adaptive Immunity to SARS-CoV-2 in Acute COVID-19 and Associations with Age and Disease Severity. Cell, 2020. 183(4): p. 996–1012 e19.

14. Tan, A.T., et al., Early induction of functional SARS-CoV-2-specific T cells associates with rapid viral clearance and mild disease in COVID-19 patients. Cell Rep, 2021. 34(6): p. 108728.

15. Rodda, L.B., et al., Functional SARS-CoV-2-Specific Immune Memory Persists after Mild COVID-19. Cell, 2021. 184(1): p. 169–183 e17.

16. Neidleman, J., et al., SARS-CoV-2-Specific T Cells Exhibit Phenotypic Features of Helper Function, Lack of Terminal Differentiation, and High Proliferation Potential. Cell Rep Med, 2020. 1(6): p. 100081.

17. Diao, B., et al., Reduction and Functional Exhaustion of T Cells in Patients With Coronavirus Disease 2019 (COVID-19). Front Immunol, 2020. 11: p. 827.

18. Tavakolpour, S., et al., Lymphopenia during the COVID-19 infection: What it shows and what can be learned. Immunol Lett, 2020. 225: p. 31–32.

19. Ziadi, A., et al., Lymphopenia in critically ill COVID-19 patients: A predictor factor of severity and mortality. Int J Lab Hematol, 2021. 43(1): p. e38–e40.

20. Dan, J.M., et al., Immunological memory to SARS-CoV-2 assessed for up to 8 months after infection. Science, 2021. 371(6529).

21. García-Abellán, J., et al., Antibody Response to SARS-CoV-2 is Associated with Long-term Clinical Outcome in Patients with COVID-19: a Longitudinal Study. J Clin Immunol, 2021. 41(7): p. 1490–1501.

22. Gong, F., et al., Peripheral CD4+ T cell subsets and antibody response in COVID-19 convalescent individuals. The Journal of Clinical Investigation, 2020. 130(12): p. 6588–6599.

23. Juno, J.A., et al., Humoral and circulating follicular helper T cell responses in recovered patients with COVID-19. Nature Medicine, 2020. 26(9): p. 1428–1434.

24. Masiá, M., et al., Durable antibody response one year after hospitalization for COVID-19: A longitudinal cohort study. J Autoimmun, 2021. 123: p. 102703.

25. Ni, L., et al., Detection of SARS-CoV-2-Specific Humoral and Cellular Immunity in COVID-19 Convalescent Individuals. Immunity, 2020. 52(6): p. 971–977 e3.

26. Robbiani, D.F., et al., Convergent antibody responses to SARS-CoV-2 in convalescent individuals. Nature, 2020. 584(7821): p. 437–442.

27. Vella, L.A., et al., T follicular helper cells in human efferent lymph retain lymphoid characteristics. J Clin Invest, 2019. 129(8): p. 3185–3200.

28. He, J., et al., Circulating precursor CCR7(lo)PD-1(hi) CXCR5(+) CD4(+) T cells indicate Tfh cell activity and promote antibody responses upon antigen reexposure. Immunity, 2013. 39(4): p. 770–81.

29. Morita, R., et al., Human blood CXCR5(+)CD4(+) T cells are counterparts of T follicular cells and contain specific subsets that differentially support antibody secretion. Immunity, 2011. 34(1): p. 108–21.

30. Brenna, E., et al., CD4(+) T Follicular Helper Cells in Human Tonsils and Blood Are Clonally Convergent but Divergent from Non-Tfh CD4(+) Cells. Cell Rep, 2020. 30(1): p. 137–152 e5.

31. Cui, D., et al., Follicular Helper T Cells in the Immunopathogenesis of SARS-CoV-2 Infection. Frontiers in Immunology, 2021. 12.

32. Thevarajan, I., et al., Breadth of concomitant immune responses prior to patient recovery: a case report of non-severe COVID-19. Nat Med, 2020. 26(4): p. 453–455.

33. Wheatley, A.K., et al., Evolution of immune responses to SARS-CoV-2 in mild-moderate COVID-19. Nat Commun, 2021. 12(1): p. 1162.

34. Locci, M., et al., Human circulating PD-1+CXCR3-CXCR5+ memory Tfh cells are highly functional and correlate with broadly neutralizing HIV antibody responses. Immunity, 2013. 39(4): p. 758–69.

35. Boppana, S., et al., SARS-CoV-2-specific circulating T follicular helper cells correlate with neutralizing antibodies and increase during early convalescence. PLoS Pathog, 2021. 17(7): p. e1009761.

36. Zhang, J., et al., Spike-specific circulating T follicular helper cell and cross-neutralizing antibody responses in COVID-19-convalescent individuals. Nat Microbiol, 2021. 6(1): p. 51–58.

37. Kestens, L., et al., Selective increase of activation antigens HLA-DR and CD38 on CD4+ CD45RO+ T lymphocytes during HIV-1 infection. Clin Exp Immunol, 1994. 95(3): p. 436–41.

38. Ziegler, S.F., F. Ramsdell, and M.R. Alderson, The activation antigen CD69. Stem Cells, 1994. 12(5): p. 456–65.

39. Townsend, L., et al., Longitudinal Analysis of COVID-19 Patients Shows Age-Associated T Cell Changes Independent of Ongoing Ill-Health. Front Immunol, 2021. 12: p. 676932.

40. Yan, Q., et al., Longitudinal Peripheral Blood Transcriptional Analysis Reveals Molecular Signatures of Disease Progression in COVID-19 Patients. J Immunol, 2021. 206(9): p. 2146–2159.

41. De Biasi, S., et al., Marked T cell activation, senescence, exhaustion and skewing towards TH17 in patients with COVID-19 pneumonia. Nature Communications, 2020. 11(1): p. 3434.

42. NIH. COVID-19 Treatment Guidelines Panel. Coronavirus Disease 2019 (COVID-19) Treatment Guidelines. National Institutes of Health. 2020 [cited 2022 9 February]; Available from: https://www.covid19treatmentguidelines.nih.gov/.

43. Mattiuzzo, G., et al., Establishment of the WHO International Standard and Reference Panel for anti-SARS-CoV-2 antibody. WHO/BS.2020.2403, 2021.

44. Jin, J.-M., et al., Gender Differences in Patients With COVID-19: Focus on Severity and Mortality. Frontiers in Public Health, 2020. 8.

45. Yang, Y., et al., Longitudinal analysis of antibody dynamics in COVID-19 convalescents reveals neutralizing responses up to 16 months after infection. Nature Microbiology, 2022.

46. Zingaropoli, M.A., et al., Major reduction of NKT cells in patients with severe COVID-19 pneumonia. Clin Immunol, 2021. 222: p. 108630.

47. Ghizlane, E.A., et al., Lymphopenia in Covid-19: A single center retrospective study of 589 cases. Annals of Medicine and Surgery, 2021. 69: p. 102816.

48. Lee, J., et al., Lymphopenia as a Biological Predictor of Outcomes in COVID-19 Patients: A Nationwide Cohort Study. Cancers (Basel), 2021. 13(3).

49. Ramljak, D., et al., Early Response of CD8+ T Cells in COVID-19 Patients. J Pers Med, 2021. 11(12).

50. He, R., et al., The clinical course and its correlated immune status in COVID-19 pneumonia. Journal of Clinical Virology, 2020. 127: p. 104361.

51. Chen, Z. and E. John Wherry, T cell responses in patients with COVID-19. Nat Rev Immunol, 2020. 20(9): p. 529–536.

52. Vella, L.A., R.S. Herati, and E.J. Wherry, CD4(+) T Cell Differentiation in Chronic Viral Infections: The Tfh Perspective. Trends Mol Med, 2017. 23(12): p. 1072–1087.

53. Crotty, S., T Follicular Helper Cell Biology: A Decade of Discovery and Diseases. Immunity, 2019. 50(5): p. 1132–1148.

54. Kervevan, J. and L.A. Chakrabarti, Role of CD4+ T Cells in the Control of Viral Infections: Recent Advances and Open Questions. Int J Mol Sci, 2021. 22(2).

55. Du, J., et al., Persistent High Percentage of HLA-DR+CD38high CD8+ T Cells Associated With Immune Disorder and Disease Severity of COVID-19. Frontiers in Immunology, 2021. 12.

56. Kared, H., et al., CD57 in human natural killer cells and T-lymphocytes. Cancer Immunol Immunother, 2016. 65(4): p. 441–52.

57. Grosso, J.F., et al., LAG-3 regulates CD8+ T cell accumulation and effector function in murine self- and tumor-tolerance systems. The Journal of Clinical Investigation, 2007. 117(11): p. 3383–3392.

58. Wolf, Y., A.C. Anderson, and V.K. Kuchroo, TIM3 comes of age as an inhibitory receptor. Nature Reviews Immunology, 2020. 20(3): p. 173–185.

59. Keir, M.E., et al., PD-1 and its ligands in tolerance and immunity. Annu Rev Immunol, 2008. 26: p. 677–704.

60. Leehan, K.M. and K.A. Koelsch, T Cell ELISPOT: For the Identification of Specific Cytokine-Secreting T Cells. Methods Mol Biol, 2015. 1312: p. 427–34.

61. Sedegah, M., The Ex Vivo IFN-γ Enzyme-Linked Immunospot (ELISpot) Assay. Methods Mol Biol, 2015. 1325: p. 197–205.

62. Larsson, M., et al., A recombinant vaccinia virus based ELISPOT assay detects high frequencies of Pol-specific CD8 T cells in HIV-1-positive individuals. AIDS, 1999. 13(7): p. 767–77.

63. Le Bert, N., et al., SARS-CoV-2-specific T cell immunity in cases of COVID-19 and SARS, and uninfected controls. Nature, 2020. 584(7821): p. 457–462.

64. Bilich, T., et al., T cell and antibody kinetics delineate SARS-CoV-2 peptides mediating long-term immune responses in COVID-19 convalescent individuals. Science Translational Medicine, 2021. 13(590): p. eabf7517.

65. Zuo, J., et al., Robust SARS-CoV-2-specific T cell immunity is maintained at 6 months following primary infection. Nature Immunology, 2021. 22(5): p. 620–626.

66. Funaro, A., et al., Role of the human CD38 molecule in B cell activation and proliferation. Tissue Antigens, 1997. 49(1): p. 7–15.

67. Glaría, E. and A.F. Valledor, Roles of CD38 in the Immune Response to Infection. Cells, 2020. 9(1).

68. Yang, X., et al., Clinical course and outcomes of critically ill patients with SARS-CoV-2 pneumonia in Wuhan, China: a single-centered, retrospective, observational study. The Lancet Respiratory Medicine, 2020. 8(5): p. 475–481.

69. Ponti, G., et al., Biomarkers associated with COVID-19 disease progression. Critical Reviews in Clinical Laboratory Sciences, 2020. 57(6): p. 389–399.

70. Akdogan, D., et al., Diagnostic and early prognostic value of serum CRP and LDH levels in patients with possible COVID-19 at the first admission. J Infect Dev Ctries, 2021. 15(6): p. 766–772.

71. Ceci, F.M., et al., Early Routine Biomarkers of SARS-CoV-2 Morbidity and Mortality: Outcomes from an Emergency Section. Diagnostics, 2022. 12(1): p. 176.

72. Enocsson, H., et al., Soluble Urokinase Plasminogen Activator Receptor (suPAR) Independently Predicts Severity and Length of Hospitalisation in Patients With COVID-19. Frontiers in Medicine, 2021. 8.

73. Fathi, N. and N. Rezaei, Lymphopenia in COVID-19: Therapeutic opportunities. Cell Biology International, 2020. 44(9): p. 1792–1797.

74. Schultheiß, C., et al., Next-Generation Sequencing of T and B Cell Receptor Repertoires from COVID-19 Patients Showed Signatures Associated with Severity of Disease. Immunity, 2020. 53(2): p. 442–455.e4.

75. Sekine, T., et al., Robust T Cell Immunity in Convalescent Individuals with Asymptomatic or Mild COVID-19. Cell, 2020. 183(1): p. 158–168 e14.

76. Wilk, A.J., et al., A single-cell atlas of the peripheral immune response in patients with severe COVID-19. Nature Medicine, 2020. 26(7): p. 1070–1076.

77. Chen, G., et al., Clinical and immunological features of severe and moderate coronavirus disease 2019. J Clin Invest, 2020. 130(5): p. 2620–2629.

78. Guan, W.-j., et al., Clinical Characteristics of Coronavirus Disease 2019 in China. New England Journal of Medicine, 2020. 382(18): p. 1708–1720.

79. Zhang, S., et al., Peripheral T cell lymphopenia in COVID-19: potential mechanisms and impact. Immunotherapy Advances, 2021. 1(1).

80. Bao, J., et al., Comparative analysis of laboratory indexes of severe and non-severe patients infected with COVID-19. Clin Chim Acta, 2020. 509: p. 180–194.

81. Qin, C., et al., Dysregulation of Immune Response in Patients With Coronavirus 2019 (COVID-19) in Wuhan, China. Clin Infect Dis, 2020. 71(15): p. 762–768.

82. Kalfaoglu, B., et al., T-Cell Hyperactivation and Paralysis in Severe COVID-19 Infection Revealed by Single-Cell Analysis. Frontiers in Immunology, 2020. 11.

83. Kundu, R., et al., Cross-reactive memory T cells associate with protection against SARS-CoV-2 infection in COVID-19 contacts. Nature Communications, 2022. 13(1): p. 80.

84. Kared, H., et al., SARS-CoV-2-specific CD8+ T cell responses in convalescent COVID-19 individuals. J Clin Invest, 2021. 131(5).

85. Cohen, K.W., et al., Longitudinal analysis shows durable and broad immune memory after SARS-CoV-2 infection with persisting antibody responses and memory B and T cells. medRxiv, 2021.

86. Cao, W.-C., et al., Disappearance of Antibodies to SARS-Associated Coronavirus after Recovery. New England Journal of Medicine, 2007. 357(11): p. 1162–1163.

87. Ng, O.-W., et al., Memory T cell responses targeting the SARS coronavirus persist up to 11 years post-infection. Vaccine, 2016. 34(17): p. 2008–2014.

88. MacLeod, M.K.L., J.W. Kappler, and P. Marrack, Memory CD4 T cells: generation, reactivation and re-assignment. Immunology, 2010. 130(1): p. 10–15.

89. Farber, D.L., N.A. Yudanin, and N.P. Restifo, Human memory T cells: generation, compartmentalization and homeostasis. Nature Reviews Immunology, 2014. 14(1): p. 24–35.

90. Tian, Y., et al., Unique phenotypes and clonal expansions of human CD4 effector memory T cells re-expressing CD45RA. Nature Communications, 2017. 8(1): p. 1473.

91. Graham, N., et al., Rapid Induction and Maintenance of Virus-Specific CD8(+) T(EMRA) and CD4(+) T(EM) Cells Following Protective Vaccination Against Dengue Virus Challenge in Humans. Front Immunol, 2020. 11: p. 479.

92. Bonifacius, A., et al., COVID-19 immune signatures reveal stable antiviral T cell function despite declining humoral responses. Immunity, 2021. 54(2): p. 340–354.e6.

93. Zenarruzabeitia, O., et al., T Cell Activation, Highly Armed Cytotoxic Cells and a Shift in Monocytes CD300 Receptors Expression Is Characteristic of Patients With Severe COVID-19. Frontiers in Immunology, 2021. 12.

94. Björkander, S., et al., SARS-CoV-2-specific B- and T-cell immunity in a population-based study of young Swedish adults. J Allergy Clin Immunol, 2022. 149(1): p. 65–75.e8.

95. Kalfaoglu, B., et al., T-cell dysregulation in COVID-19. Biochem Biophys Res Commun, 2021. 538: p. 204–210.

96. Kotredes, K.P., B. Thomas, and A.M. Gamero, The Protective Role of Type I Interferons in the Gastrointestinal Tract. Frontiers in Immunology, 2017. 8.

97. Belz, G.T., J.D. Altman, and P.C. Doherty, Characteristics of virus-specific CD8(+) T cells in the liver during the control and resolution phases of influenza pneumonia. Proceedings of the National Academy of Sciences, 1998. 95(23): p. 13812–13817.

98. Saeidi, A., et al., T-Cell Exhaustion in Chronic Infections: Reversing the State of Exhaustion and Reinvigorating Optimal Protective Immune Responses. Frontiers in Immunology, 2018. 9.

99. Jurek, T., et al., SARS-CoV-2 Viral RNA Is Detected in the Bone Marrow in Post-Mortem Samples Using RT-LAMP. Diagnostics, 2022. 12(2): p. 515.

100. Han, X., et al., Six-month Follow-up Chest CT Findings after Severe COVID-19 Pneumonia. Radiology, 2021. 299(1): p. E177–e186.

101. Zhang, S., et al., Eight months follow-up study on pulmonary function, lung radiographic, and related physiological characteristics in COVID-19 survivors. Scientific Reports, 2021. 11(1): p. 13854.

102. Zhang, P., et al., Long-term consequences in lung and bone associated with hospital-acquired severe acute respiratory syndrome: a 15-year follow-up from a prospective cohort study. The Lancet, 2018. 392: p. S11.

103. Okoye, I.S., et al., Coinhibitory Receptor Expression and Immune Checkpoint Blockade: Maintaining a Balance in CD8(+) T Cell Responses to Chronic Viral Infections and Cancer. Front Immunol, 2017. 8: p. 1215.

104. Wu, H., et al., Postmortem high-dimensional immune profiling of severe COVID-19 patients reveals distinct patterns of immunosuppression and immunoactivation. Nature Communications, 2022. 13(1): p. 269.

105. Long, Q.X., et al., Clinical and immunological assessment of asymptomatic SARS-CoV-2 infections. Nat Med, 2020. 26(8): p. 1200–1204.

106. Ibarrondo, F.J., et al., Rapid Decay of Anti–SARS-CoV-2 Antibodies in Persons with Mild Covid-19. New England Journal of Medicine, 2020. 383(11): p. 1085–1087.

107. Sherina, N., et al., Persistence of SARS-CoV-2-specific B and T cell responses in convalescent COVID-19 patients 6–8 months after the infection. Med, 2021. 2(3): p. 281–295.e4.

108. Aleem, A., A.B. Akbar Samad, and A.K. Slenker, Emerging Variants of SARS-CoV-2 And Novel Therapeutics Against Coronavirus (COVID-19), in StatPearls. 2022, StatPearls Publishing, Copyright © 2022, StatPearls Publishing LLC.: Treasure Island (FL).

109. Carfì, A., et al., Persistent Symptoms in Patients After Acute COVID-19. JAMA, 2020. 324(6): p. 603–605.

110. Huang, C., et al., 6-month consequences of COVID-19 in patients discharged from hospital: a cohort study. The Lancet, 2021. 397(10270): p. 220–232.

111. Xie, Y., et al., Long-term cardiovascular outcomes of COVID-19. Nature Medicine, 2022.

