## Supplementary Tables and Figures for "Long-term T cell perturbations and waning antibody levels in individuals needing hospitalization for COVID-19"

### Supplementary figures and tables Govender et al

**Supplementary Table 1.**
**Monoclonal antibodies used for quantification of lymphocytes in whole blood**

| Marker | Fluorochrome | Company | Clone |
|---|---|---|---|
| **CD57** | FITC | BD | HNK-1 |
| **IgD** | FITC | BD | IA6-2 |
| **CD27** | PE | BD | L128 |
| **CD45** | PerCP | BD | 2DI |
| **CD38** | PerCP-Cy5.5 | BD | HIT2 |
| **CD19** | PE-Cy7 | BD | SJ25C1 |
| **CD56** | PE-Cy7 | BD | NCAM16.2 |
| **CD8** | APC | BD | SK1 |
| **CD21** | APC | BD | B-ly4 |
| **CD20** | APC-H7 | BD | L27 |
| **CD4** | Horizon V450 | BD | RPA-T4 |
| **CD3** | Horizon V500 | BD | UCHT1 |

Antibodies used at 4µl to 50ul whole blood

**Supplementary Table 2.**
**Antibodies used for T cell characterization**

| Marker | Fluorochrome | Company | Clone | Concentration |
|---|---|---|---|---|
| **CD3** | BV750 | BD | SK7 | 1/160 |
| **CD45** | APC | BD | HI30 | 1/400 |
| **CD4** | Spark NIR 685 | BioLegend | SK3 | 1/80 |
| **CD8** | AF700 | BioLegend | SK1 | 1/400 |
| **HLA-DR** | BV570 | BioLegend | L243 | 1/40 |
| **CD38** | APC Fire 810 | BioLegend | HIT2 | 1/40 |
| **CD57** | Pacific blue | BioLegend | HCD57 | 1/400 |
| **PD1** | PE-Cy5.5 | Novus Biologicals | J116 | 1/40 |
| **CD45RA** | FITC | BD | HI100 | 1/200 |
| **CCR7** | PE | BioLegend | G043H7 | 1/20 |
| **CD27** | PE-Cy7 | BioLegend | O323 | 1/40 |
| **CD28** | PerCP | ThermoFisher | CD28.2 | 1/20 |
| **CD25** | BV650 | BD | M-A251 | 1/40 |
| **CD127** | BV785 | BioLegend | A019D5 | 1/40 |
| **CD95** | BV605 | BioLegend | DX2 | 1/40 |
| **CXCR5** | BV510 | BD | RF8B2 | 1/20 |
| **TIM-3** | BV421 | BioLegend | F38-2E2 | 1/80 |
| **LAG-3** | PE-CF594 | BD | T47-530 | 1/40 |
| **ICOS** | BV711 | BD | DX29 | 1/40 |
| **CCR6** | PerCP-eFlour 710 | ThermoFisher | R6H1 | 1/80 |
| **CXCR3** | BV480 | BD | 1C6 | 1/20 |
| **CD69** | APC-H7 | BD | FN50 | 1/40 |
| **Viability** | ViaKrome 808 | Beckman Coulter | | 1/50 |

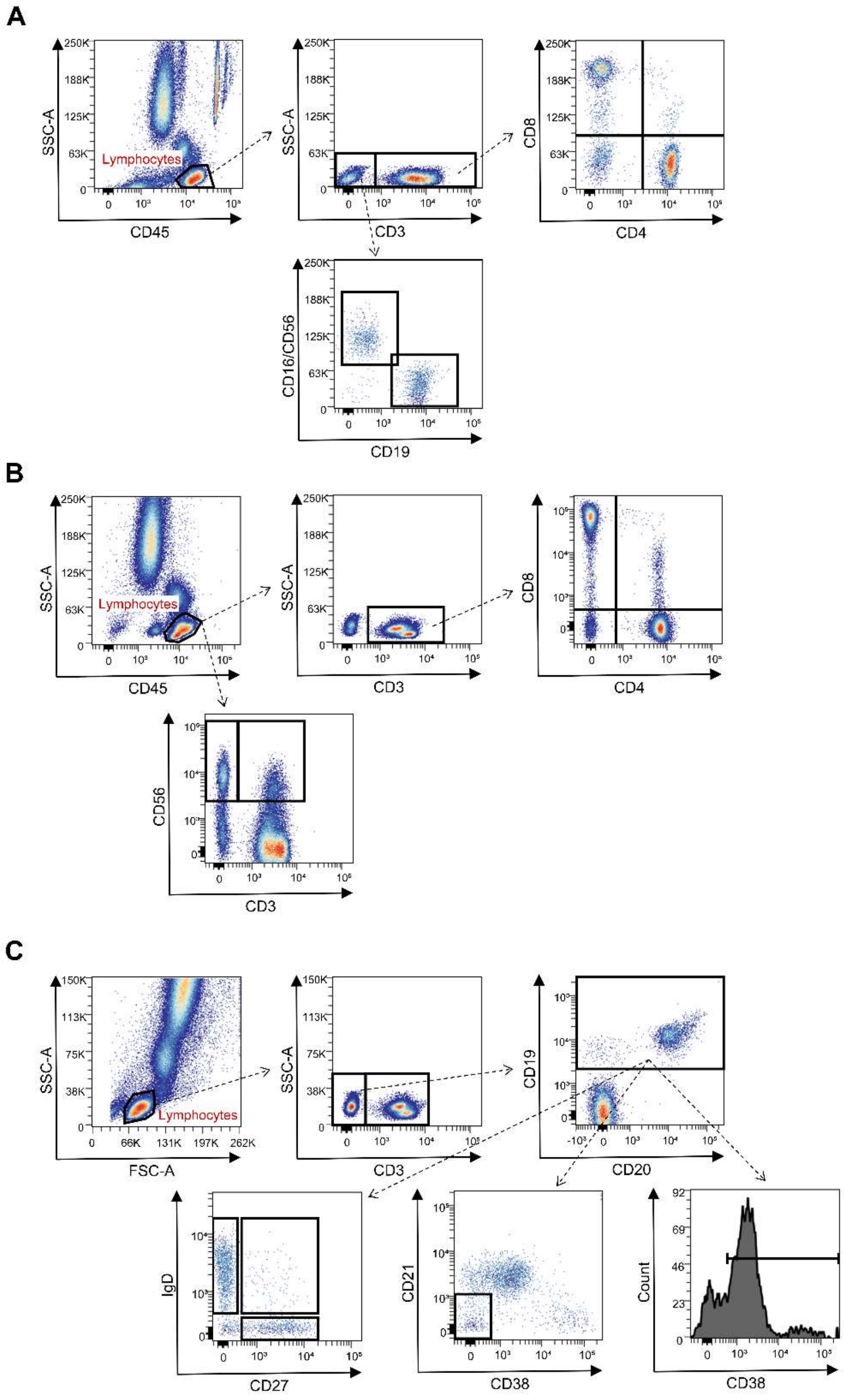


**Supplementary Figure 1.**

**Gating strategy to define immune cell populations at inclusion**

Representative gating strategy to analyze the cellular clinical parameters. (**A**) Representative gating for quantification of B cell and T cell populations quantities using truecount tubes with conjugated CD45, CD3, CD4, CD8, CD16/CD56 monoclonal antibodies. (**B**) Representative gating to determine proportions of T cells and NK cells using CD45, CD3, CD4, CD8, CD56. (**C**) Representative gating to determine B cell proportions and activation states using CD3, CD19, CD20, IgD, CD27, CD21, CD38.

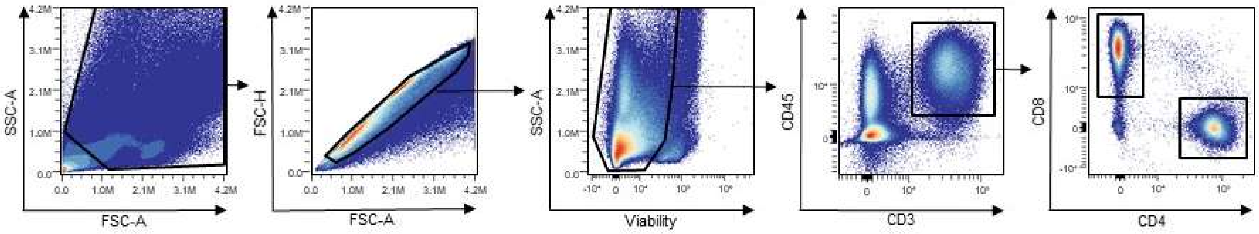


**Supplementary Figure 2.**

**Gating strategy to define bulk and CD4+ and CD8+ T cells.**

Suspensions of PBMCs were prepared by blocking and antibody staining for analysis by spectral flow cytometry. Representative gating strategy is shown here, depicting how $CD4^{+}$ and CD8+ lymphocytes were selected for further analysis, using ViaKrome808 viability dye, CD3, CD45, CD4 and CD8 monoclonal antibodies.

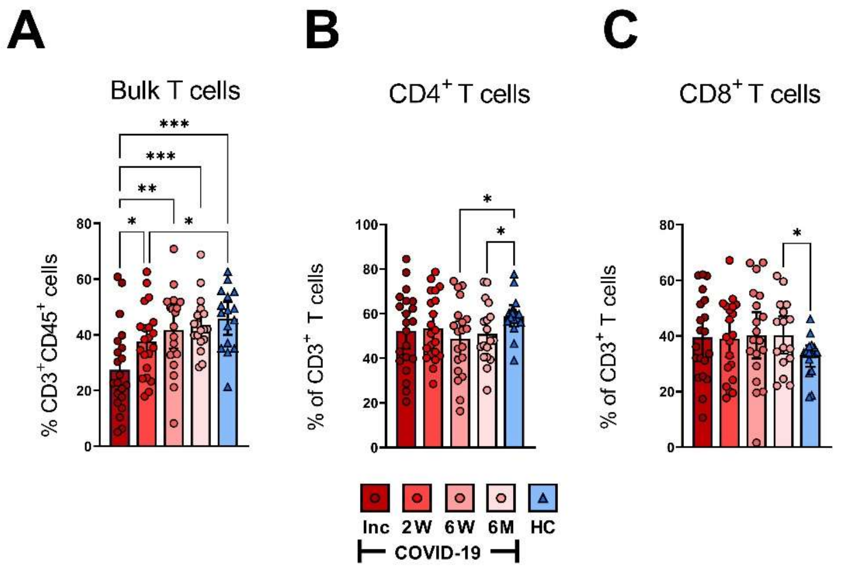


**Supplementary Figure 3.**

**Bulk, CD4⁺ and CD8⁺ T cells during COVID-19 infection.**

PBMCs obtained from 23 COVID-19 patients and 16 healthy donors over a 6-7 month period were stained with CD45, CD3, CD4 and CD8 monoclonal antibodies and assessed by flow cytometry to phenotype the major T cells subsets: (**A**) $CD3^+$, (**B**) $CD4^+$ T cells and (**C**) $CD8^+$ T cells, as shown in percentage. Data is represented as mean with 95% Cl, with significance of *$p \leq 0.05$, **$p \leq 0.01$, ***$p \leq 0.001$, ****$p \leq 0.0001$, determined using Brown-Forsythe and Welch ANOVA tests. Inc = Inclusion in study at the hospital, 2W = 2 weeks, 6W = 6 weeks, 6M = 6-7 months, HC = healthy control.

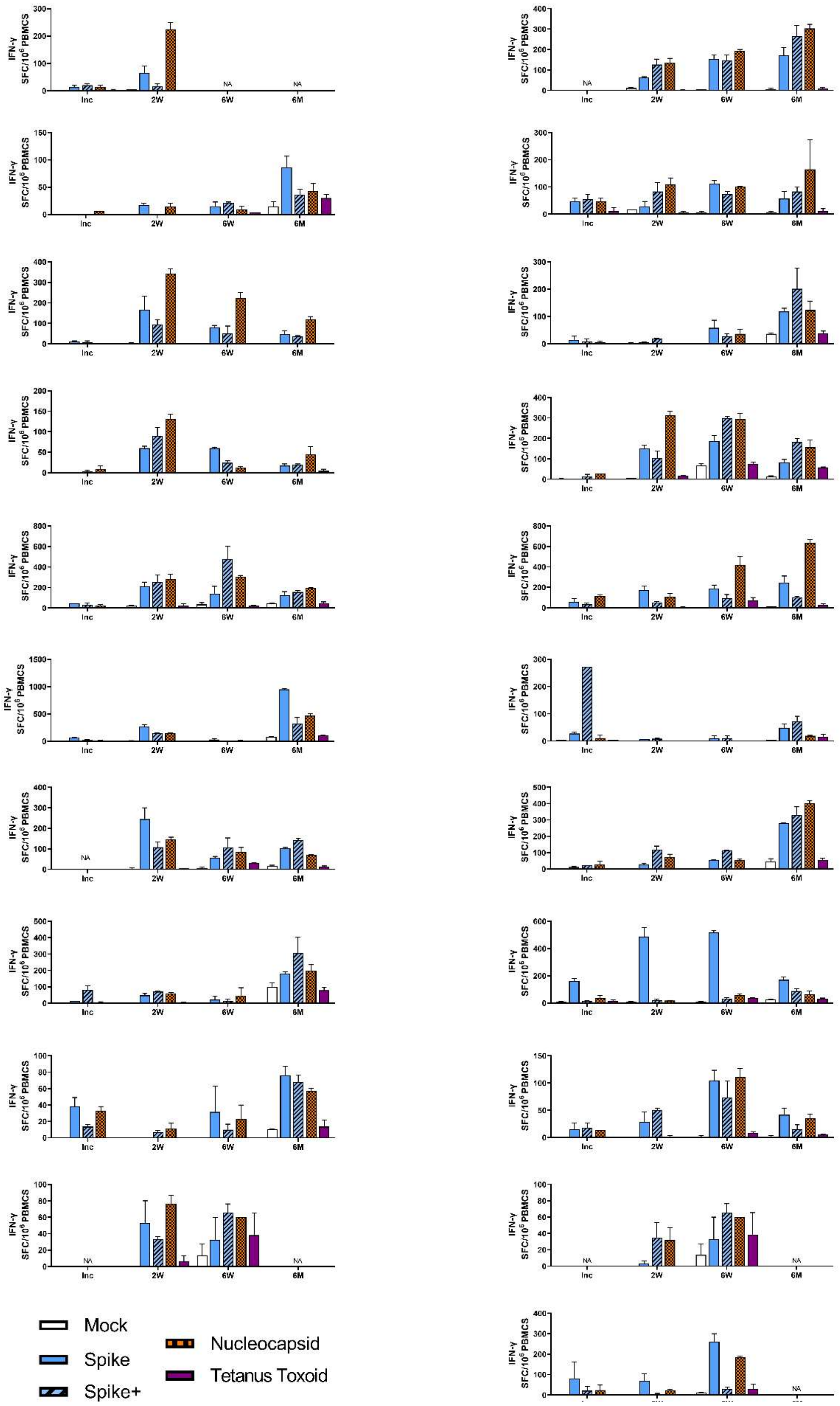


**Supplementary Figure 4.**

**SARS-CoV-2-specific T cells in individual COVID-19 patients.**

PBMCs collected from 21 covid patients were unstimulated (Mock) or stimulated with SARS-CoV-2 specific overlapping peptides from: Spike, spike+, and nucleocapsid and tetanus toxoid protein. The IFN-γ production was measured after 48h as spot forming cells (SFC) by ELISPOT. Each graph represents a single donor. Data is represented as mean with 95% CI, with significance of $**p<0.01$, $***p < 0.001$, $***p < 0.0001$ determined using Brown-Forsythe and Welch ANOVA tests.
